# Gating of Memory to Behavior by the Locus Coeruleus

**DOI:** 10.1101/2024.01.09.574947

**Authors:** Tianyu Wang, Xinyang Zhang, Haoyu Duan, Dan Xia, Tianxiang Li, Rongzhen Yan, Yang Zhan, Yulong Li, Wen-Jun Gao, Qiang Zhou

## Abstract

An essential function of memory is to guide behavior for survival and adaptation. While considerable knowledge has been accumulated on memory formation, much less is understood about how retrieved memories direct behavior/action. In the auditory Pavolovian threat conditioning paradigm, retrieval of conditioned threat memory activates dorsomedial prefrontal (dmPFC) neurons exhibiting transient responses (T-neurons), which activate both dmPFC neurons exhibiting sustained responses (S-neurons) and locus coeruleus (LC) neurons. Auditory inputs to S-neurons enable the conversion from transient to sustained responses so that the freezing durations match those of the auditory cues. Activation of LC neurons is required for the conversion by enhancing S-neuron responses, which, interestingly, opens a short time window during which non-conditioned cues also lead to freezing. The transition from memory to behavior thus hinges on the integration of retrieved memory, sensory inputs, and emotional/body state cues to generate a selective, adequate, and finely tuned behavior.

**Significance statement:** This study provides new insights into the neural circuitry and mechanisms of how retrieved memories direct the execution of behavior in response to conditioned threatening stimuli. It reveals how different neuron types in the dmPFC interact with LC neurons to determine and modulate the duration and intensity of defensive responses. It also shows that activation of LC neurons can induce the generalization of freezing to non-threatening cues, which may have implications for understanding anxiety disorders. This study contributes to the field of neuroscience by advancing the understanding of memory-behavior conversion and role of the dmPFC and LC in conditioned threat/fear behavior.

## Introduction

How memory is converted into motor outputs to drive an adequate behavior remains a long-standing and important question in neuroscience. Memory allows the association to form between a cue predicting significant events with appropriate responding behavior patterns for higher chances of survival and adaptation (Pavlov 1927; Rescorla, 1988; Pearce and Bouton, 2001; Fanselow et al., 2015). Much effort has been devoted to elucidating the mechanisms underlying memory formation and modulation, while how a formed memory elicits behavior is far less understood. Given that behavior itself is the ultimate determinant of survival and fitness, it is important to gain a deeper understanding of the transition from memory to behavior and the modulation of this process.

Pavlovian/classical conditioning has been widely used as a model to study the learning and memory process due to its simplicity and robustness (Rescorla, 1988; LeDoux, 2000; Maren and Quirk, 2004; Fanselow et al., 2015). In this model, the conditioned stimulus (CS) mimics the detection of a predator, which triggers a range of post-encounter responses (Fanselow, 1989). One commonly used conditional response is freezing, which is a defensive response characterized by the cessation of movement and a state of immobility, and it reduces the likelihood of being detected and attacked by a predator (Fanselow and Lester, 1988). In auditory threat conditioning, the coincident occurrence of a CS (sound) and an unconditioned stimulus (US, footshock) leads to an association between CS and US, which is encoded in the threat memory (LeDoux, 2000; Maren and Quirk, 2004; Fanselow et al., 2015; Tovote et al., 2015; Herry and Johansen, 2014). Reencountering the same CS activates the formed threat memory and elicits a set of behavioral responses, indicating that a memory directs behavior. More specifically, threatening conditioning results in synaptic potentiation in the LA/basolateral amygdala (LA/B) and enhanced spiking in these neurons (Maren and Quirk, 2004; Hans-Christian and Pare, 2010).

Reactivating this threat memory in LA/B neurons initiates a cascade of events, including the activation of dorsomedial prefrontal cortex (dmPFC) neurons, followed by activation of neurons in BLA and central amygdala (CeA), and ultimately the expression of freezing behavior (LeDoux, 2000; Sotres-Bayon and Quirk, 2010; Herry and Johansen, 2014; Gilmartin et al., 2014; Duvarci and Pare, 2014; Tovote et al., 2015). We have used the above circuit as a model system to examine how memory transits to behavior.

Previous studies have shown that the dmPFC acts as a hub to integrate sensory and contextual information to modulate the behavioral responses elicited by the retrieved emotional memory (Sotres-Bayon and Quirk, 2010; Tovote et al., 2015; Herry and Johansen, 2014). During retrieval of auditory threat memory with a long CS (a few to dozens sec), LA/B neurons displayed transient spiking (∼ 1 sec) (Quirk et al., 1995; Paréand Collins., 2000; Goosens and Maren, 2004), while similar transient responses were also seen in a population of dmPFC neurons (Gilmartin et al., 2005; Courtin et al., 2014; Yan et al., 2019, 2021). Another dmPFC population of excitatory neurons displayed sustained responses matching the duration of CS in a manner similar to the elicited freezing behavior (Burgos-Robles et al., 2009; Pendyam et al., 2013). These dmPFC neurons with sustained responses are known to project to BLA and, hence, likely direct behavioral responses (Burgos-Robles et al., 2009). These findings suggest the participation of two distinct dmPFC neuronal populations in transitioning from the amygdala-stored conditioned memory to defensive/freezing behavior.

This hypothesized division of labor between the dmPFC transient (T-) and sustained (S-) neurons predicts the distinct inputs, outputs, and properties of these two dmPFC populations. Specifically, the T-neurons are presumably suited for memory processing by integrating relevant inputs and emotional states to determine whether a memory is expressed or suppressed. One major modulator of the T-neuron activity is the dmPFC PV-neurons which integrate various inputs and act to suppress the formed memory/learned responses (Courtin et al., 2014; Yan et al., 2019, 2021; Wang et al., 2023). In contrast, S-neuron responses may match the CS duration and respond to neuromodulators, such as norepinephrine (Burgos-Robles et al., 2009; Pendyam et al., 2013). Thus, S-neurons are likely influenced by the emotional/bodily states during memory retrieval. However, it remains unknown whether S-neurons are activated by T-neurons, a representation of threat memory in PFC. Furthermore, S-neurons are likely driven by sensory inputs (such as sound) based on their CS-locked responses, but this hypothesis has not been tested.

Norepinephrine (NE) is released by locus coeruleus (LC) neurons when an organism enters a fearful state, and NE modulates cortical function during vigilance, attention, arousal, and stress (Aston-Jones et al., 1991; Berridge et al., 1993; Berridge and Waterhouse, 2003; Morilak et al., 2005; Sara, 2009; Arnsten, 2009). It is well-established that NE profoundly affects the formation and extinction of fear/threat memory (Uematsu et al., 2017; Giustino and Maren, 2018; Likhtik E and Johansen, 2019). In addition, NE has been shown to sustain PFC neuronal spiking and freezing in response to threat CS (Rodriguez-Romaguera et al., 2009; Pendyam et al., 2013). However, the contribution of LC/NE to the transition from memory to behavior remains unknown. For example, what are the inputs to the LC neurons that trigger this NE release? Does LC/NE exert the classical amplification effect as shown for the sensory system? Does NE modify the retrieved memory itself or only the associated behavior? These are essential questions that can be addressed using the auditory threat conditioning model to better understand LC/NE functions in general.

Here we demonstrated that: (1) dmPFC T-neurons are memory units and ensure that memory retrieval is selective to the conditioned cues; (2) dmPFC S-neurons are behavior/action units that convert transient responses to sustained ones to match the cue inputs; (3) LC neurons, which are selectively activated by PFC T-neuron inputs, enhance the S-neurons responses. Interestingly, once activated, further activation of LC neurons is not selective to the conditioned cues. Thus, a successful transition from memory to behavior requires the activation of stored memory, continued inputs from the sensory area, and the concomitant activation of LC neurons.

## Materials and Methods

### Animals

Male C57BL6/J were purchased from Guangdong Medical Laboratory Animal Center (China). Male C57BL6/J were group housed (5 - 6 mice/cage) under a 12 hr light/dark cycle (8:00 AM to 8:00 PM) and were provided with food and water *ad libitum*. Mice of 3 - 5 months of age were used. For *in vivo* electrophysiology experiments, mice were singly housed before the surgery to implant multi-wire electrodes. They were habituated by gentle handling for 3 days before behavioral and recording experiments. Bedding, water, and food were changed every week. All animal experiments were performed in accordance with the ARRIVE (Animal Research: Reporting of *In Vivo* Experiments) guidelines, approved by the Peking University Shenzhen Graduate School.

### In vivo recording

Mice of 3-5 months of age were used. Mice were anesthetized with isoflurane (induction 3%, maintenance 1.5%). Two screws were implanted to secure electrode array implants. Multiwire electrodes were unilaterally implanted in the targeted brain regions with coordinates: dmPFC (1.98 mm anterior to Bregma; ± 0.3 mm lateral to midline, and 1.1–1.3 mm below cortical surface); LA/B (1.75 mm posterior to Bregma; ± 3.3 mm lateral to midline and 3.65 mm below cortical surface); BLA (−1.31 mm posterior to Bregma, ± 3.05 mm lateral to midline and 4.75 mm below cortical surface). Electrodes consisted of 16 individually insulated nichrome wires (35μm inner diameter, impedance 300 ∼ 900 KΩ; Stablohm 675, California Fine Wire) arranged in a 3×5×5×3 pattern (∼ 200 μm spacing between wires). Electrodes were attached to an 18-pin connector (Mil-Max) and secured with dental cement. Optrode arrays are composed of an optic fiber (250 μm) and multiwire electrodes (the end of the optical fiber is ∼ 200 - 300 μm above the tips of the recording electrodes).

After surgery, mice were allowed to recover for 7 - 14 d. Broadband (0.3 Hz to 7.5 kHz) neural signals were simultaneously recorded (16 bits at 30 kHz) from implanted 16-channel arrays using a 64-channel data acquisition system (Zeus, Bio-Signal Technologies). At completion, recorded extracellular spikes were aligned and sorted (offline sorter software, Plexon), and further analyzed using NeuroExplorer (Nex Technologies) and MATLAB (Yan et al., 2019). Responses during CS were normalized to pre-tone spike rate by calculating a Ζ-score (Yan et al., 2019). For statistical analysis, comparisons were performed using the averaged Ζ-score values calculated during the 500 ms period after CS onset.

Neurons were considered CS responsive if they showed significant, time-locked changes in their spiking upon CS presentation. Neuron spiking within ™1 to 0 s as an average basal level, when spiking above the average level adds 3×S.E.M will be used as CS responsive.

### In vivo imaging

To record Ca^2+^ responses in the LC neurons, mice were injected with the AAV2/9-CaMKII-GCaMP7s virus in the LC. To record NE responses in the PFC and LA/B, mice were injected with the rAAV-hSyn-NE2h-WPRE-hGH polyA virus in the PFC and LA/B. The procedure for fiber photometry has been described (Yan et al., 2019). Data were analyzed using MATLAB. Changes in fluorescence values (ΔF/F) were calculated as (F-F0)/F0, where F0 is the baseline fluorescence averaged between ™10 and ™2 s before CS presentation. The magnitudes of responses were calculated by measuring the area under the curve (AUC) in a period of 2 s from the CS onset.

### Slice recording and opto-stimulation

For projection-specific activation, mice were injected with AAV-CamKII-cre virus and Retrobeads bilaterally in the LA/B and AAV-DIO-ChR2 virus in the dmPFC to activate LA/B inputs. Slicing and recording procedures were similar to that used in Yao et al., 2018. Briefly, to record spontaneous excitatory post-synaptic currents (sEPSCs) in S-neurons, somatic whole-cell voltage clamp recordings (−60 mV) were obtained from layer II/III red excitatory neurons in dmPFC. To record T-neurons spontaneous excitatory post-synaptic currents (sEPSCs), somatic whole-cell voltage clamp recordings (−60 mV) were obtained from layer II/III green excitatory neurons in dmPFC. To confirm a direct projection from T-neurons to S-neurons, we injected CaMKII-cre anterior transport virus in LA/B and DIO-ChR2-eYFP virus in dmPFC, and injected retrobeads (red) in BLA to label dmPFC-projecting LA/B neurons. The dmPFC T-neurons were of eYFP (green) and S-neurons of retrobeads (red). To record neurons’ spontaneous inhibitory post-synaptic currents (sIPSCs), somatic whole-cell voltage clamp recordings (+5 mV) were obtained from layer II/III excitatory neurons in dmPFC. Current clamp recordings were used to examine spikes evoked by a series of 500 ms depolarizing current pulses with 4 s intervals, and each step with an increase of 25 pA (from 0 to 375 pA). All neurons were recorded for at least 5 minutes. Resting membrane potentials (RMPs) were measured under I = 0 pA condition, and neurons with RMPs above ™50 mV were excluded from the analysis.

### Behavioral assays

Threat conditioning and retrieval test took place in two different chambers/contexts (for details, see Wang et al., 2023). Scoring freezing behavior was performed using a video recording system (Coulbourn Instruments). Mice were considered freezing if no movement was detected for at least 2 s. On day 0, mice were habituated in context A and received 4 CS (conditioned stimulus; 30 s, 60 dB, 50-ms pips tone (3 kHz). Threat conditioning was conducted on day 1, with CS co-terminated with a US (unconditioned stimulus; 2 s foot shock, 0.75 mA), for 3 CS-US pairing trials. Training trials were separated by 90 s intertrial intervals. For threat memory retrieval on day 2, conditioned mice received 4 trials of various combinations of CS+ and CS- (30 s, 60 dB; CS+: 50 ms pips tone (3 kHz); CS-: white noise) in context B at 24 hr after conditioning, including 30 s CS+, 2 s CS+/28 s CS- or 2 s blue light stimulation/28 s CS- in context B at 24 hr post-conditioning. For experiments with NE sensor imaging, conditioned mice received 3 trials of 30 s CS+, 10 s CS+, 2 s CS+, 2 s CS+/28 s CS- or 2 s blue light stimulation/28 s CS- in context B at 24 hr post-conditioning.

### In vivo optogenetic manipulations

The C57 mice were injected with AAV2/9-CaMKII-eNpHR3.0-mCherry or AAV2/9-CaMKII-ChR2-eYFP virus bilaterally in the LA/B and opto-fiber placed in the dmPFC to inhibit or activate LA/B-projecting dmPFC excitatory neurons (Fig. 3A, 3D, 3E and 7C). C57 mice were injected with AAV2/9-Ef1α-cre (retro) virus bilaterally in the BLA and AAV2/9-Ef1α-DIO-eNpHR3.0-mCherry viruses in dmPFC to inhibit dmPFC-projecting BLA neurons (Fig. 3B). To inhibit LA/B projecting dmPFC neurons, 593-nm laser was used (30 s, constant light, ∼ 10 mW at end of the optical fibers). To inhibit dmPFC-projecting LA/B neurons, a 593-nm laser was used (30 s, constant light, ∼ 10 mW at the end of the optical fibers). To activate LA/B- dmPFC projections, a 473-nm laser was used (2 s or 1 s, 20 ms pulse width, 20 Hz, ∼ 10 - 15 mW). C57 mice were injected with AAV2/9- Ef1α-cre (retro) virus in the dmPFC, AAV2/9-Ef1α-DIO-eNpHR3.0-mCherry viruses in LC and opto-fiber placed in the LC to inhibit LC-projecting dmPFC neurons (Fig. 6G). C57 mice were injected with AAV2/9-CaMKII-cre virus in LA/B, AAV2/9-Ef1α-DIO-ChR2-mCherry viruses in dmPFC and optical fiber placed in the dmPFC to activate LA/B-projecting dmPFC T-neuron (Fig. 7F). C57 mice were injected with AAV2/9-CaMKII-ChR2-mCherry virus in the LC, and optical fiber was placed in the LC to activate LC excitatory neurons (Fig. 7H).

### Cell harvesting

The T- and S-neurons were identified in brain slices using their selective fluorescence markers, respectively. Weak negative pressure was applied to aspirate the entire neuron into the glass patch electrodes, which were treated with DEPC water (ThermoFisher) in advance. The pipette tip was broken onto the wall of a 0.2 ml tight-lock tube (TubeOne) to allow the entire neuron to be immersed in a 1 μL drop of RNase-free lysis buffer (provided by Beijing Genomics institution (BGI)) placed on the side of the tube. The tube was kept on ice and two neurons were collected in one tube within 5 min. The tube was then placed in dry ice until 20 neurons were harvested from each mouse. Samples were rapidly spun down (5 - 10 s) and stored at - 80°C before reverse transcription. Reverse transcription, PCR amplification and sequencing were performed by BGI.

### RNA Sequencing

A total of 6 samples were sequenced separately and were performed as before (Xia et al., 2022). The expression amount of identified RNAs was analyzed using TPM (transcripts per million). Differential expression analysis was conducted using DESeq2 package in R software (1.2.22). Differentially expressed genes (DEGs) were defined as those with |fold change| ≥ 2 and q-value (a corrected P-value using Benjamini and Hochberg multiple testing correction) < 0.05.

### Statistical analysis and Graphing

Figures were generated using Dr. Tom online system and Adobe Illustrator CC 2018. Statistical significance was calculated using a two-tailed paired/unpaired t Test, one-way/two-way repeated measures (RM) ANOVA (version 8, GraphPad Prism Software), as noted. Data are reported as mean ± SEM. Significance levels are noted as *, *p*<0.05; **, *p*<0.01; ***, *p*<0.001.

## Results

### Transient and sustained neuronal responses in PFC associated with conditioned threat cues

By using previously established methods and criteria, we simultaneously examined spiking in the dmPFC neurons and freezing in mice (Yan et al., 2019; 2021; Wang et al., 2023). Although both transient and sustained neuronal responses have been shown in the PrL/dmPFC neurons after auditory threat conditioning, they have not been demonstrated in the same mice. Thus, we first recorded dmPFC neuron responses to CS presentation in conditioned mice showing significant levels of freezing during CS presentation (Fig. 1A). Indeed, CS-elicited two spiking patterns in two non-overlapping neuronal populations recorded in dmPFC, with either transient (T-neurons) (Fig. 1B) or sustained (S-neurons) responses (Fig. 1C). As reported previously, T-neuron responses last about 1 sec (Fig. 1B) while the duration of S-neuron responses roughly matched the duration of CS (Fig. 1C). Both increased and decreased sustained responses were observed, but only increased transient responses were seen (Sup Fig. 1). There were no significant responses to CS-, indicating the selectivity of the responses to conditioned stimulus (Sup Fig. 1). Out of the 692 dmPFC neurons recorded, 22% (151/692) showed transient while 6% (41/692) showed sustained responses, 6% (40/692) were inhibitory neurons with increased response (based on their spike waveforms, Sup Fig. 2) (Fig. 1D). The rest of the recorded dmPFC neurons did not display significant CS-induced changes in spiking. For the remainder of the work, we only presented the results from dmPFC excitatory neurons that respond to the CS.

**Figure 1.**
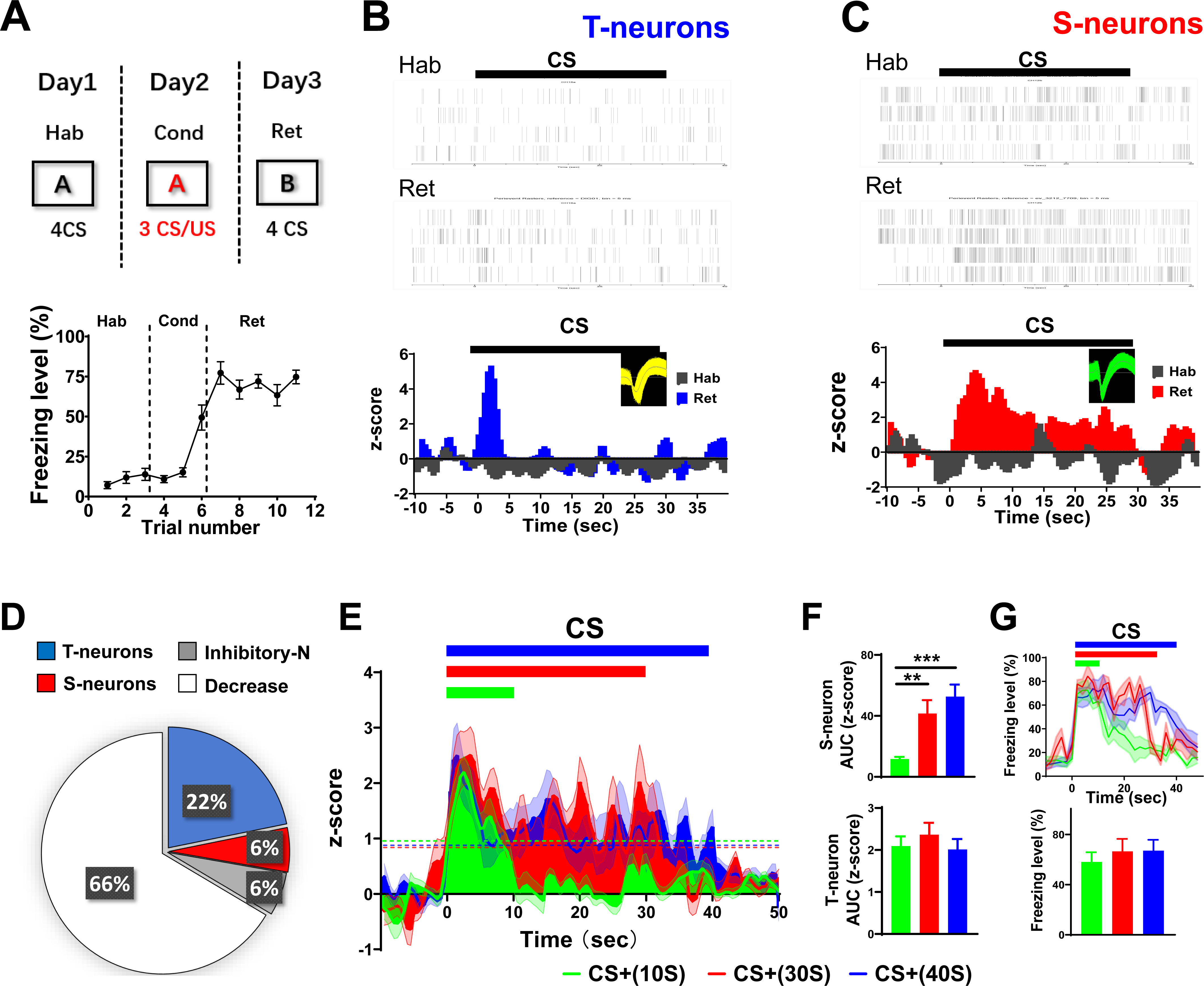
Activity of the dmPFC T-neurons and S-neurons during retrieval of threat memory. (A) (Upper) Experimental procedure for threat conditioning. (Lower) Percent freezing before, during, and after auditory threat conditioning. N= 12 mice. (B) (Upper) Raster plots of spiking in representative T-neurons during habituation or CS retrieval after conditioning. (Lower) Perievent histograms showing spiking responses of T-neurons. Bars indicate CS presentation. Bin width, 0.5 s. n=23 units/5 mice. (C) (Upper) Raster plots of spiking in representative S-neurons during habituation or CS retrieval after conditioning. (Lower) Perievent histograms of S-neuron responses. n=17 units/5 mice. (D) Distribution of dmPFC neurons based on their responses to CS after conditioning. Of the recorded neurons, 22% (151/692) showed a transient increase, 6% (41/692) sustained increase; 6% (40/692) were inhibitory neurons and 66% (460/692) showed no response. n= 692 units/42 mice. (E) Spiking responses in S-neurons elicited by CSs with different durations. (F) (Upper) The AUC (z-score) of CS-elicited S-neuron response. One-way RM ANOVA, F (2, 45) = 9.671, Bonferroni’s posttest; CS+ (10 s) vs. CS+ (30 s), P < 0.01; CS+ (10 s) vs. CS+ (40 s), P < 0.001; n= 16 units/8 mice. (Lower) The AUC (z- score) of CS-elicited responses in T-neurons (n= 29 units/8 mice). (G) (Upper) Freezing durations elicited by three CSs (One-way RM ANOVA, F (2, 18) = 15.98, Bonferroni’s posttest; CS+ (10 s) vs. CS+ (30 s), P < 0.01; CS+ (10 s) vs. CS+ (40 s), P < 0.001). (Lower) Freezing level (%) elicited by three CSs (N= 5 mice). Unless specified, statistical comparisons were performed using two-tailed unpaired t tests; ∗, P < 0.05; ∗∗, P < 0.01; ∗∗∗, P < 0.001. Data represented as mean ± SEM.

In our experiments, a 30-s long CS was used during both the conditioning and retrieval tests. To determine whether the durations of conditioned freezing responses and S-neuron spiking match that of CSs with different durations, we used three different CS durations (10, 30 and 40 s). We observed the durations of S-neuron responses were well-matched with the corresponding CSs (Fig. 1E, 1F), while T-neurons responded with similar amplitude regardless of CSs tested (Fig. 1F). In addition, the durations of freezing matched the CS durations with freezing levels comparable (Fig. 1G). These results suggest that neurons contributing to the conditioned threat responses likely receive direct information about CS (such as duration). In contrast, regardless of the specific CS used, T-neuron responses were of the same magnitude and duration. In comparison, the majority of LA/B excitatory neurons only showed transient responses (Sup Fig. 3), consistent with prior observations (Quirk et al., 1995; Pare and Collins, 2000; Goosens and Maren, 2004).

### Distinct encoding of threat responses by T-neurons and S-neurons

We have shown previously that the T-neuron responses are directly related to freezing level under a few different conditions, including response to CS representing safety or low probability of CS-US association (Yan et al., 2019; 2021; Wang et al., 2023). However, S-neuron responses under these conditions are unknown. We thus plotted threat responses (freezing level) against neuronal responses (spike frequency) in three conditioning paradigms: 100% CS-US association (100P), 50% CS-US association (50P), and safety learning (SL). Here, 100% CS-US association refers to 1CS:1US pairing, while 50% CS-US association refers to 2CS:1US pairing, and SL is defined as CS- (safety cue) unpaired with US/foot shock, which resulted in reduced responses to safety cue. We first generated averaged neuronal responses for each mouse, and found a significant correlation between freezing and averaged neuronal responses with transient (1 s) or sustained (30 s) responses (Fig. 2A). This finding suggests that when taking the entire neuronal population into account, both transient and sustained responses match well with behavioral responses. We then plotted the total spiking of all responding neurons (i.e., summed responses in T-neurons or S-neurons) against freezing levels for each mouse. We observed a significant correlation between freezing levels and responses in T-neurons or S-neurons (Fig. 2B), suggesting that the responding neurons alone are sufficient for effectively encoding CS-induced threat responses.

**Figure 2.**
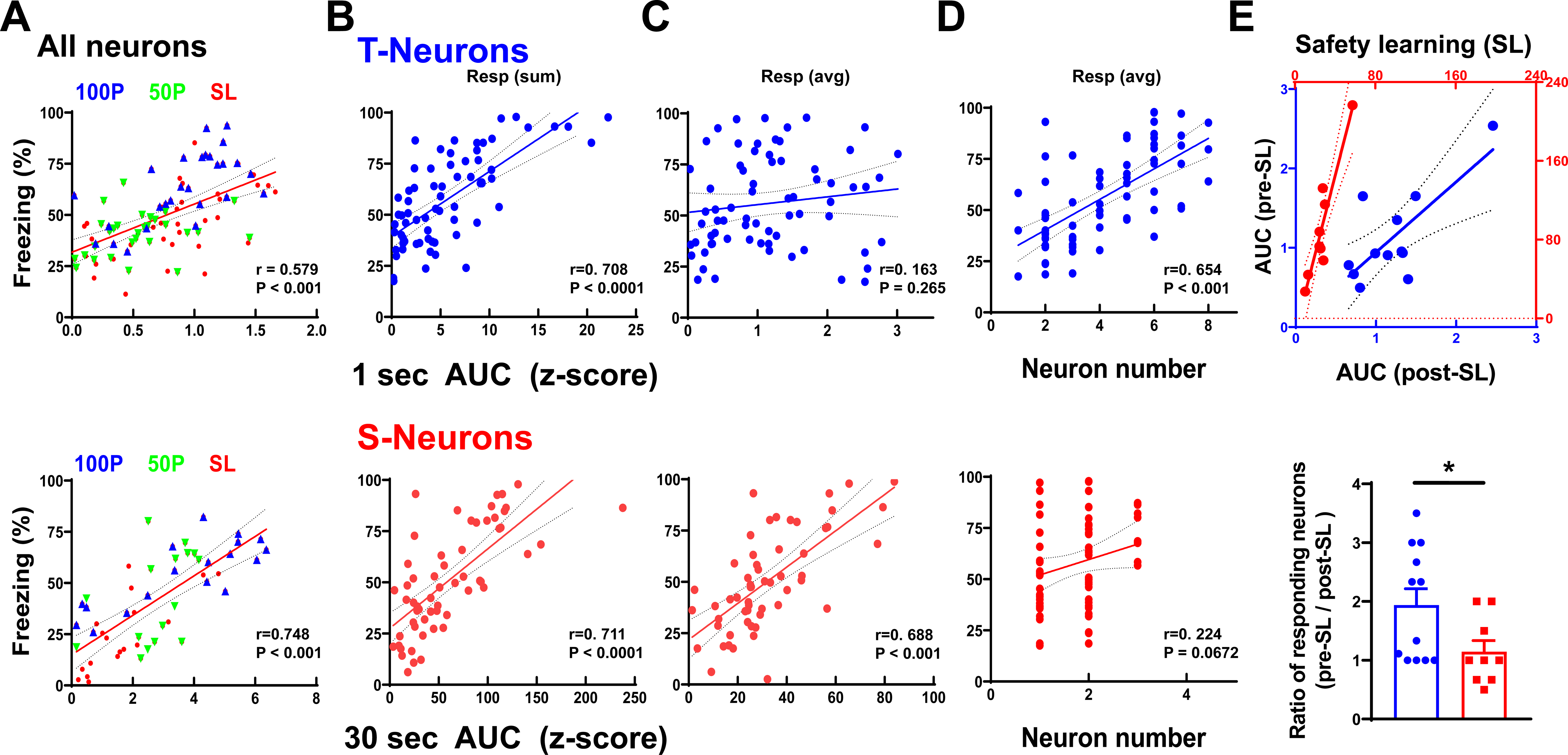
Distinct encoding of conditioned threat responses by T-neurons and S- neurons. (A) Linear regression analysis between freezing levels and spiking during the first second of CS (top) or the entire CS (30 s, bottom) in all dmPFC neurons. Note that the sustained responses only included those neurons with sustained spiking. Responses were calculated as AUC (area under the curve) and averaged AUC for each mouse was plotted. Each symbol represents one mouse. Data from conditioning with 100% CS-US association (100P), 50% association (50P), and safety learning (SL) groups were included. Linear regression, T-neuron, r = 0.579, R^2^ = 0.335, P < 0.001; S-neuron, r = 0.748, R^2^ = 0.56, P < 0.001; 50P, N= 28 mice; 100P, N= 29 mice; safety learning, N= 12 mice. (B) Linear regression analysis between freezing level and spiking responses in T- or S-neurons from responding neurons. Total AUC from all included neurons in a given mouse was plotted. Each symbol represents one mouse. Data from 100P and 50P conditioning groups. Linear regression, T-neurons, r = 0.708, R^2^ = 0.502, P < 0.001; S-neurons, r = 0.711, R^2^ = 0.506, P < 0.001; N= 69 mice. (C) Linear regression analysis between freezing level and spiking responses in T- or S-neurons. Averaged AUC for all recorded T- or S-neurons in a given mouse was used. Each symbol represents one mouse. Data from 100P and 50P conditioning groups. Linear regression, T-neurons, r = 0.163, R^2^ = 0.0185, P = 0.265; S-neurons, r = 0.688, R^2^ = 0.474, P < 0.001; N= 69 mice. (D) Relationship between freezing level and number of T- or S-neurons in a given mouse. Data from 100P and 50P conditioning groups. Linear regression, T-neurons, r = 0.654, R2 = 0.428, P < 0.001; S-neurons, r = 0.225, R2 = 0.051, P = 0.067; N= 69 mice. (E) (Top) CS-elicited neuronal responses (AUC before safety learning/AUC after safety learning) for T-neurons or S-neurons. Data from safety learning group. Two-tailed paired t-test, t = 3.091, df = 11, S-neurons, pre-SL vs. post-SL, P < 0.05; N= 12 mice. (Bottom) Ratio of responding neurons before over after SL (Pre-SL/Post-SL) for T-neurons or S-neurons. Data from safety learning group. Two-tailed unpaired t test, t = 2.202, df = 19, P < 0.05; N= 12 mice.

To understand whether individual T-neurons or S-neurons function in a similar manner, we plotted the averaged responses of all T-neurons or S-neurons for each mouse. Here we found a big difference between T-neurons and S-neurons: a significant correlation between freezing level and S-neurons but not for T-neurons (Fig. 2C). Hence, S-neurons likely encode threat responses in a graded manner based on the responses of individual neurons, whereas T-neurons may employ population encoding with each neuron responding in an all-or-none manner. The above results predict that the number of responding neurons likely plays different roles for T- and S-neurons. When the numbers of responding T- or S-neurons were plotted against freezing levels, we found an excellent relationship for T-neurons but not for S-neurons (Fig. 2D). Put together, the above results indicate two distinct encoding mechanisms: population encoding for T-neurons and magnitude encoding for S-neurons.

The above conclusion was obtained from a comparison between different groups of animals. To avoid potential influence/bias in this population-based analysis, we tested the above encoding model in the same mouse under different threat states. We took advantage of a high freezing level to CS- during the fear generalization state and a low freezing level after safety learning (SL; *i.e.*, after CS- has become the safety cue) (Yan et al., 2019). We found that the response magnitudes in S-neurons were significantly reduced after safety learning compared to that before learning. In contrast, the responses in T-neurons were maintained at a similar level (Fig. 2E).

Regarding the number of responding neurons, we observed that the ratio of pre-SL over post-SL was mostly above 1 for T-neurons, while this ratio is about 1 for S-neurons (Fig. 2E), consistent with S-neurons using response magnitude while T-neurons using population encoding mechanism. This reduced magnitude of S-neuron responses after SL is likely caused by the reduced numbers of responding T-neurons. These results provide further evidence for different encoding mechanisms for T- and S-neurons.

### Major connections of T-neurons and S-neurons

The differential contributions of T- and S-neurons to the threat responses suggest that their connection patterns and intrinsic neuronal properties may also differ. To examine whether this is the case, we first examined their afferents, efferents, and connections with each other. Previous studies showed that T-neurons receive synaptic inputs from LA/B (LeDoux, 2000; Quirk et al., 1995; Yan et al., 2019, 2021), and we thus tagged the LA/B inputs to dmPFC using injection of CaMKII-NpHR-mCherry virus in LA/B. In the same set of mice, opto-inhibition of the LA/B terminals in the dmPFC during CS retrieval significantly reduced freezing level and spiking in both T-neurons and S-neurons (Fig. 3A). Since it is known that dmPFC neurons project to basolateral amygdala (BLA), we tagged the BLA-projecting dmPFC neurons using retrograde virus (retro-cre virus in dmPFC and DIO-NpHR in BLA). Opto-inhibition of these BLA-projecting dmPFC neurons significantly reduced freezing level (Fig. 3B). Importantly, a significantly lower spiking was seen in the S-neurons but not T-neurons (Fig. 3B). These results indicate that CS-induced signals transmit from LA/B to dmPFC and to BLA, and S-neurons are downstream of T-neurons. Supporting this conclusion, spiking in the S-neurons lagged spiking of T-neurons by approximately 200 msec during CS retrieval (Fig. 3C). During opto-stimulation of the LA/B inputs, we observed the same 200 msec delay between spikes in T-neurons and S-neurons, with T-neuron leading (Fig. 3D). Interestingly, during opto-stimulation the S-neuron responses were transient, suggesting that their sustained spiking is caused by the long CS.

**Figure 3.**
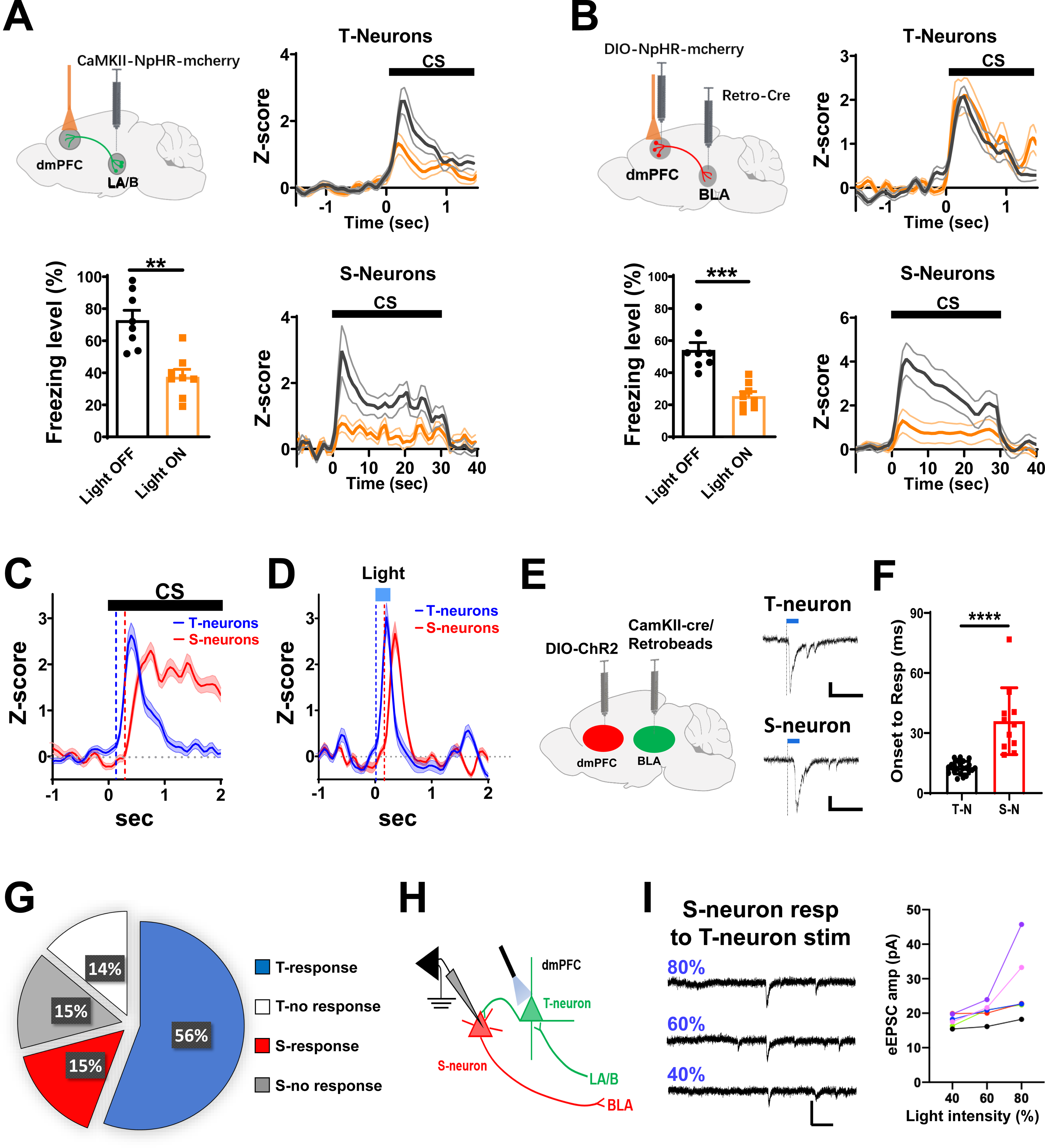
Main inputs and outputs of T-neurons and S-neurons, and connections between them. (A) (Left) Opto-inhibition of LA/B-dmPFC projections on freezing levels during CS retrieval. NpHR3.0 viral infection with yellow light illumination. Two-tailed paired t- test, t = 3.891, df = 7, Laser ON vs. Laser OFF, P < 0.01; N= 8 mice. (Right) Opto-inhibition of LA/B-dmPFC projections on spike rates in T-neurons and S-neurons. T- neurons, n= 35 units/8 mice; S-neurons, n= 20 units/8 mice. (B) (Left) Opto-inhibition of BLA-projecting dmPFC neurons on freezing levels during CS retrieval. Two-tailed paired t test, t = 5.679, df = 7, Light ON vs. Light OFF, P < 0.01; N= 8 mice. (Right) Opto-inhibition of BLA-projecting dmPFC neurons during CS retrieval on spike rates in dmPFC T-neurons or S-neurons. T- neurons, n= 41 units/8 mice; S-neurons, n= 24 units/8 mice. (C) Latency of CS-elicited spiking in T-neurons and S-neurons in threat-conditioned mice. Vertical lines represent the slope (dy/dx) of responses equal to 1 (the points of intersection were 0.122 and 0.298 sec, respectively.). T-neurons, n= 54 units/12 mice; S-neurons, n= 29 units/12 mice. (D) Latency to spike in T-neurons and S-neurons to opto-activation of LA/B-dmPFC projections (bar). Vertical lines represented the slope (dy/dx) of responses equal to 1 (the points of intersection were 0.008 and 0.159 sec, respectively). T-neurons, n= 32 units/6 mice; S-neurons, n= 16 units/6 mice. (E) (Left) Diagram showing the virus injection for examining the responses in T- neurons and S-neurons to opto-activation of LA/B inputs in PFC slices. (Right) Representative traces of EPSCs recorded in T- or S-neurons to opto-stimulation (bar) in PFC slices. Vertical lines represent the start of stimulation. Scale bars, 100 ms and 10 pA. (F) Latency of responses to opto-activation of LA/B inputs in dmPFC T-neurons or S- neurons in PFC slices. Two-tailed unpaired t-test, t = 6.783, df = 26, T-neuron vs. S- neuron, P < 0.0001; T-neurons, n= 19 cells/5 mice; S-neurons, n= 9 cells/4 mice. (G) Percentage of PFC neurons to opto-activation of LA/B inputs in PFC slices. Of the recorded neurons, 56% (44/79) of T-neurons showed responses and 14% (12/79) showed no response; 15% (12/79) of S-neurons showed responses and 15% (12/79) showed no response. (H) Schematic diagram showing opto-activation of T-neurons while recording in S- neurons in PFC slices. (I) (Left) Representative traces of recorded EPSCs in a single S-neuron to opto-stimulation of T-neurons with increasing light power. (Right) Population responses. n= 5 cells. Scale bars, 200 ms and 40 pA.

To obtain more definitive results on the connections between T- and S-neurons, we used combined whole-cell patch recording and opto-stimulation in acute PFC slices. Here, T-neurons were identified by their responses to LA/B inputs, while S- neurons were by their BLA projections (see Methods). First, we found that delays in response onset to LA/B opto-stimulation were significantly longer in S-neurons compared to T-neurons (Fig. 3E, 3F). Second, to confirm a direct projection from T- neurons to S-neurons, we infected CaMKII-positive excitatory neurons in LA/B with CaMKII-cre anterior transport virus and DIO-ChR2-eYFP virus in dmPFC, and injected retrobeads (red) in BLA to label dmPFC-projecting LA/B neurons. This strategy effectively labeled dmPFC T-neurons with eYFP (green) and S-neurons with retrobeads (red), and enabled optogenetic stimulation of the T-neurons that express ChR2. We recorded PFC neurons by opto-activation of LA/B inputs in the PFC slices. Of all recorded neurons, we found that 56% (44/79) of T-neurons showed responses and 14% (11/79) showed no response. In addition, 15% (12/79) of S-neurons showed responses, and 15% (12/79) had no response (Fig. 3G). Moreover, the majority of S- neurons responded to opto-stimulation of T-neurons (9/12 S-neurons tested, 75%), indicating direct activation of S-neurons by T-neurons (Fig. 3H). A few S-neurons showed a step-like increase in their responses with a maximum of 3 steps (Fig. 3I). This result suggests that a single S-neuron may be connected to as many as 3 T- neurons. Collectively, the above results strongly indicate that T-neurons are the first station receiving LA/B inputs in PFC and they project to S-neurons.

### Distinct neuronal properties of T-neurons and S-neurons

Our results thus far suggest that T- and S-neurons are two distinct excitatory neuronal populations co-existing in the dmPFC. To understand whether they may possess distinct neuronal and synaptic properties, we first conducted an RNAseq analysis. We aspirated RNAs from the labeled T-neurons or S-neurons from brain slices using patch electrodes and examined their RNA profile (Xia et al., 2022). A total of 14609 genes were detected (Table S1), with 650 showing significantly higher expression and 728 with lower expression in the S-neurons compared to T-neurons (Fig. 4A and Table S1). Principal components analysis of 6 samples allowed distinction between T-neurons and S-neurons (Fig. 4B). We found that: (1) for ion channels, 8 genes (*Kctd4, Kctd10, Kctd12, Kcne3, Kcng2, Kcna5, Kcnt2 and Kcnd2*) were significantly higher and 4 genes (*Kctd18, Kcnab3, Kcna4 and Kcns1*) were significantly lower, in the S-neurons (Fig. 4C and Table S2), with *Cacnb2* significantly higher in the S-neurons, and with no difference in the sodium channel genes (Table S2). These results suggest differences in neuronal excitability between T-neurons and S-neurons. (2) No difference was observed for GABA receptors and glutamate receptors between T-neurons and S-neurons, except for *Gabrr2* and *Grik1* which were lower and higher in the S-neurons, respectively (Table S2). (3) Higher expression of *Adrb1* (adrenergic receptor, β-1) in the S-neurons than T-neurons (Fig. 4D and Table S2). (4) T-neurons and S-neurons showed distinct cell-adhesion molecule (CAM) expression profiles, suggesting different cell-surface assembly codes (Fig. 4E and Table S2). Taken together, the above RNA expression profile supports T-neurons and S-neurons as two distinct neuronal populations.

**Figure 4.**
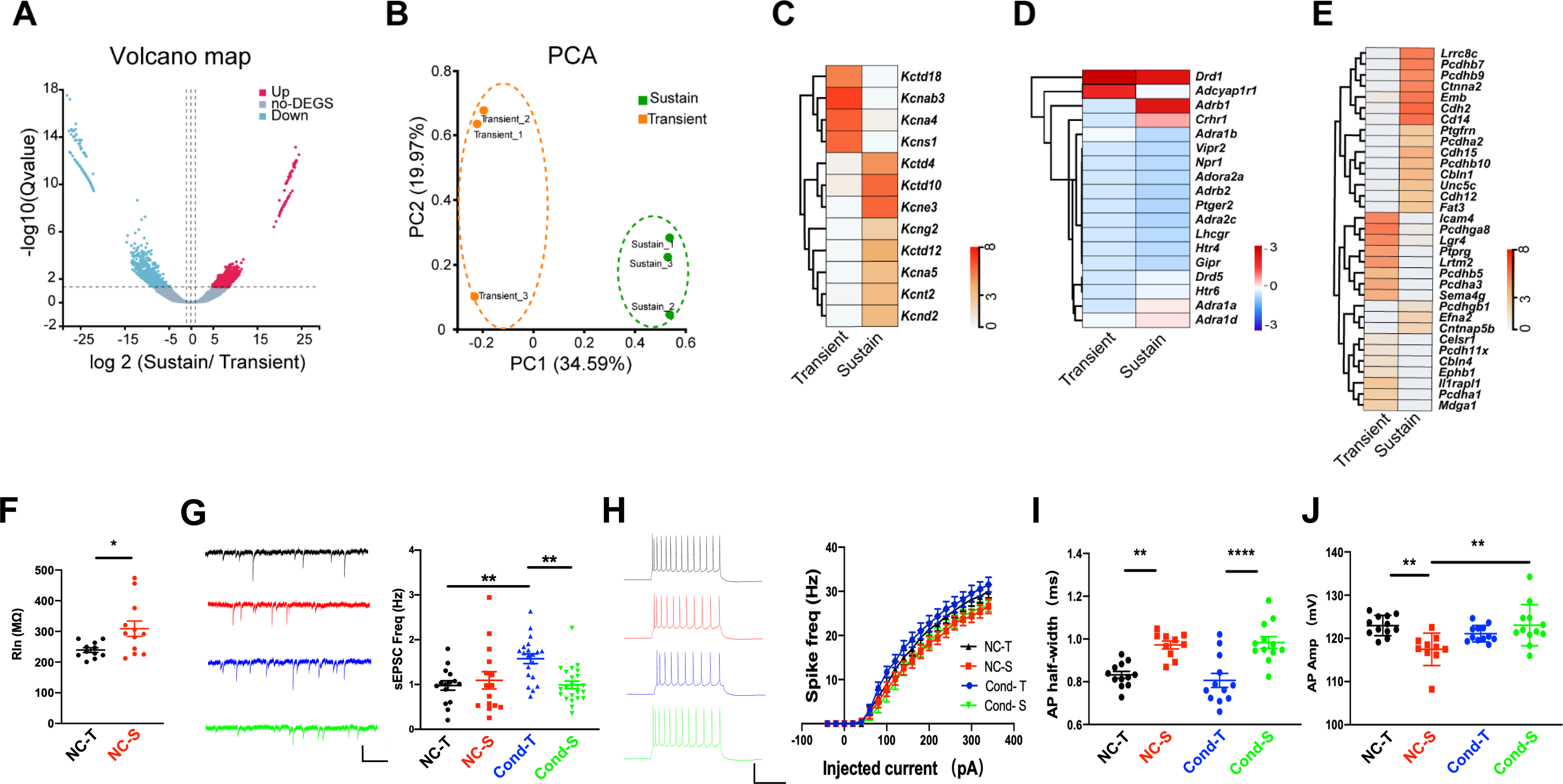
Characterizations of T-neurons and S-neurons. (A) Volcano plots showing significantly differentially expressed genes (DEGs) between T-neurons and S-neurons. (B) Principal components analysis (PCA) of an RNA-seq data set of 6 samples was sufficient to distinguish between T-neurons and S-neurons. (C) Heat map showing average transcripts per million (TPM) of significantly differentially expressed genes encoding potassium channels. (D) Heat map showing average TPM of significantly differentially expressed genes related to norepinephrine. Heatmap was standardized by rows. (E) Heat map showing average TPM of significantly differentially expressed genes encoding cell adhesion molecules. In all heatmaps, the redder the color, the higher the expression level. Standardized method: log (average TPM+1). n= 20 cells/mouse, 3 mice in each group. (F) (Left) Representative sEPSC traces and (right) sEPSC frequency in dmPFC T- neurons and S-neurons. One-way RM ANOVA, F (2,53) = 7.804, Bonferroni’s posttest, NC-T vs. Cond-T, P < 0.01; NC-S vs. Cond-S, P < 0.01; n= 15 cells/4 mice (NC-T), 15 cells/4 mice (NC-S), 20 cells/4 mice (Cond-T), 24 cells/6 mice (Cond-S). Scale bars, 20 pA and 500 ms. (G) Input resistance in T-neurons and S-neurons. Two-tailed unpaired t-test, NC-T vs. NC-S, P = 0.0179; n= 11 cells/3 mice (NC-T), 13 cells/3 mice (NC-S). NC, non-conditioned. (H) (Left) Representative traces of action potentials in dmPFC T-neurons and S-neurons elicited by injection of current through the recording electrodes. (Right) Spike frequency of action potentials plotted against injected currents. Two-way RM ANOVA, F (30,990) = 6.894, Bonferroni’s posttest, P < 0.001; NC-T vs. NC-S, P < 0.01; NC-T vs. Cond-T, P < 0.001; n= 10 cells/4 mice (NC-T), 11 cells/4 mice (NC- S), 11 cells/4 mice (Cond-T), 21 cells/6 mice (Cond-S). Scale bars, 50 mV and 200 ms. (I) Half-width of action potentials in T-neurons and S-neurons. One-way RM ANOVA, F (3, 42) = 13.57, P < 0.001, Bonferroni’s posttest; NC-T vs. NC-S, P < 0.01; Cond-T vs. Cond-T, P < 0.001; n= 12 cells/4 mice (NC-T), 10 cells/4 mice (NC- S), 12 cells/4 mice (Cond-T), 12 cells/6 mice (Cond-S). (J) Amplitude of action potentials in T-neurons and S-neurons. One-way RM ANOVA, F (3, 41) = 5.530, P < 0.01, Bonferroni’s posttest; NC-T vs. Cond-T, P < 0.05; n= 10 cells/4 mice (NC-T), 9 cells/4 mice (NC-S), 12 cells/4 mice (Cond-T), 14 cells/6 mice (Cond-S).

We then recorded from T-neurons and S-neurons in the dmPFC slices and compared their electrophysiological properties. We found significantly higher input resistance in the S-neurons (Fig. 4F). Excitatory inputs to these neurons, measured using sEPSCs, were not different between T-neurons and S-neurons (Fig. 4G; Sup Fig. 4). Interestingly, S-neurons showed a conditioning-dependent increase in sEPSC frequency (Fig. 4G), likely caused by higher release probability, since sEPSC amplitude was unaltered (Sup. Fig. 4A). The intrinsic excitability was not different between T-neurons and S-neurons (Fig. 4H). The half-width of spikes was significantly longer in the S-neurons (Fig. 4I), while spike amplitudes were significantly lower in the S-neurons (Fig. 4J; Sup Fig. 4F), and other properties of spikes were not different (Sup. Fig. 4B-E). Spike amplitude in the S-neurons also showed a conditioning-dependent increase, consistent with S-neurons being more sensitive to modification by experience.

### Sustained S-neuron responses and freezing require TeA inputs and PFC NE elevation

The observation that S-neuron responses match well with the CS durations (Fig. 1E) suggests that S-neurons are influenced by sensory (sound) inputs. Previous studies showed that the temporal association cortex (TeA) mediates the specific type of sound used in our experiments (Cambiaghi et al., 2015; Cho et al., 2016; Kwon et al., 2012). To examine whether TeA represents the primary input to dmPFC, we first injected retrobeads into the dmPFC and observed that the majority of beads were present in the TeA but not the primary auditory cortex (Fig. 5A). Optogenetic stimulation of TeA inputs elicited significant responses in the identified S-neurons in the dmPFC slices (Fig. 5B). Both observations support significant TeA inputs to the dmPFC S-neurons.

**Figure 5.**
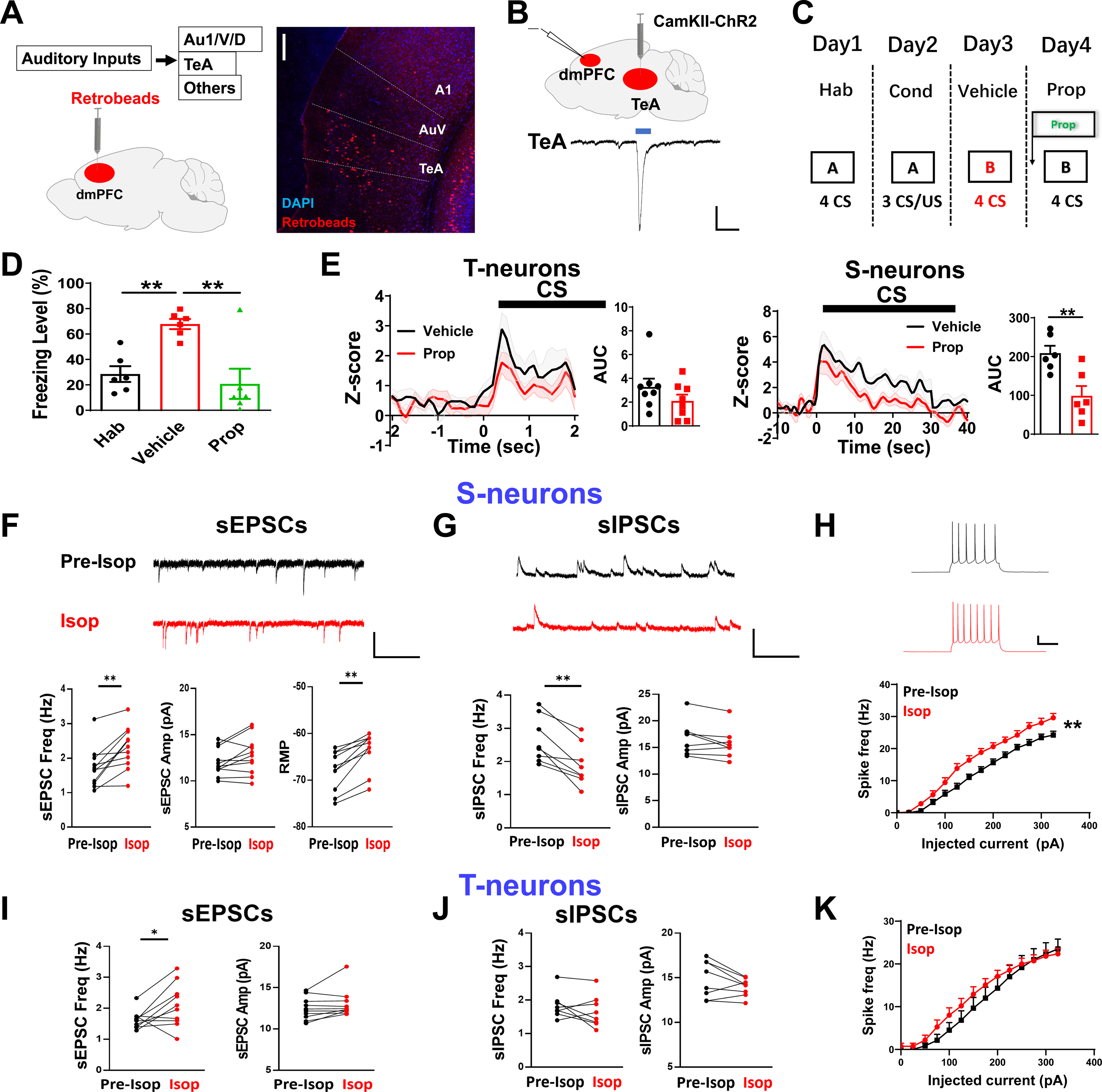
Contribution of β-norepinephrine receptors to dmPFC neuronal responses and freezing. (A) Experimental paradigm for examining inputs to dmPFC using an injection of retrobeads in dmPFC (left) and locations of the retrobeads in the auditory cortex (right). A1, primary auditory cortex; AuV, auditory cortex; TeA, temporal association cortex. Scale bar, 200 μm. (B) EPSCs in dmPFC S-neurons to opto-stimulation of TeA inputs, with CaMKII- ChR2 virus expressed in TeA. Bar indicates blue light illumination. Scale bars, 200 pA and 50 ms. (C) Experimental procedure for examining dmPFC injection of propranolol on the neuronal responses and freezing levels in conditioned mice. (D) Freezing levels during habituation (Hab), retrieval after conditioning with (Ret Prop) or without (Ret) propranolol (prop) injection. Two-tailed paired t-test, t = 9.869, df = 5, Hab vs. Ret, P < 0.01; N= 6 mice. Two-tailed paired t test, t =3.463, df = 5, Ret vs. Ret prop, P < 0.01; N= 6 mice. (E) Effect of propranolol injection in PFC on spike rates in T-neurons (left) and S- neurons (right). Two-tailed paired t-test, t = 3.53, df = 10, Ret vs. Ret Prop, P < 0.01; T-neurons, n= 10 units/6 mice; S-neurons, n= 11 units/6 mice. (F) (Upper) Bath application of Isop (50 μM) on sEPSCs in S-neurons. Scale bars, 20 pA and 50 m. (Lower) Effects of Isop on sEPSC frequency (Two-tailed paired t-test, t = 4.246, df = 10, Pre-Isop vs. Isop, P < 0.01; n= 11 cells/4 mice), sEPSC amplitude (Two-tailed paired t-test, t = 0.976, df = 20, Pre-Isop vs. Isop, P > 0.05; n= 11 cells/4 mice) and resting membrane potential (RMP) in S-neurons (Two-tailed paired t-test, t = 6.143, df = 9, Pre-Isop vs. Isop, P < 0.01; n= 10 cells/4 mice). (G) (Upper) Bath application of Isop (50 μM) on sIPSCs in S-neurons. Scale bars, 50 pA and 50 m. (Lower) Effects of Isop on sIPSC frequency (Two-tailed paired t-test, t = 4.961, df = 7, Pre-Isop vs. Isop, P < 0.01; n= 8 cells/3 mice) and sIPSC amplitude in S-neurons (Two-tailed paired t-test, t = 2.088, df = 7, Pre-Isop vs. Isop, P > 0.05; n= 8 cells/3 mice). (H) (Upper) Sample traces of action potentials in S-neurons before and after bath application of Isop. Scale bars, 20 mV and 200 ms. (Lower) Effects of Isop on the intrinsic excitability of S-neurons. Two-way RM ANOVA, F (15, 288) = 2.294, Bonferroni’s posttest, P < 0.01; Pre-Isop vs. Isop, P < 0.01; n= 12 cells/3 mice (Pre-Isop), n= 12 cells/3 mice (Isop). (I) Effects of Isop on sEPSC frequency (Two-tailed paired t-test, t = 2.333, df = 9, Pre-Isop vs. Isop, P < 0.05; n= 10 cells/4 mice) and sEPSC amplitude (Two-tailed paired t-test, t = 1.758, df = 9, Pre-Isop vs. Isop, P < 0.05; n= 10 cells/4 mice) in T- neurons. (J) Effects of Isop on sIPSC frequency (Two-tailed paired t test, t=1.265, df=7, Pre-Isop vs. Isop, P > 0.05; n= 8 cells/3 mice) and sIPSC amplitude (Two-tailed paired t- test, t = 1.778, df = 7, Pre-Isop vs. Isop, P > 0.05; n= 8 cells/3 mice) in T-neurons. (K) Effects of Isop on the intrinsic excitability of T-neurons. Two-way RM ANOVA, F (15, 336) = 0.4195, Bonferroni’s posttest, P > 0.05; Pre-Isop vs. Isop, P > 0.05; n= 12 cells/3 mice.

Neuromodulators have a critical influence on encoding and expression of memory. Of particular relevance to threat conditioning, signaling by NE via β- adrenergic receptors has been shown to maintain S-neuron responses and freezing (Burgos-Robles et al., 2009; Pendyam et al., 2013). However, whether this action is mediated by β-adrenergic receptors in the PFC is unclear. We first performed a local infusion of β-noradrenergic antagonist propranolol (100 nmol/mL) into the dmPFC (Fig. 5C) which resulted in a significant reduction in both freezing levels (Fig. 5D) and S-neuron responses (Fig. 5E) during CS retrieval test. Importantly, propranolol did not significantly affect the spike rate in T-neurons (Fig. 5E), indicating the NE impact is downstream of T-neuron activation.

NE is known to enhance the activity/excitability of postsynaptic neurons and neurotransmitter release. Our RNAseq results also suggest a higher expression of β- adrenergic receptors in the S-neurons (Fig. 4D). Hence, we tested whether S-neurons may show preferentially higher responses to NE than the T-neurons. To do so, we bath applied β-AR agonist isoprenaline (ISOP, 50 μM), which led to a significant increase in the sEPSC frequency but not amplitude (Fig. 5F) in the PFC slices, suggesting an increased presynaptic glutamate release onto the S-neurons. ISOP induced a significant depolarization of the resting membrane potentials in the S- neurons (Fig. 5F) and a significant reduction in the sIPSC frequency but not amplitude (Fig. 5G). A significant increase in the intrinsic excitability by ISOP was also observed in the S-neurons (Fig. 5H). In contrast, the main impact of ISOP on the T-neurons was an increased sEPSC frequency (Fig. 5I-5K; Sup Fig. 5). These results indicate a preferential effect of NE on the S-neurons to enhance their activity via higher excitatory inputs, lower inhibitory inputs, and higher intrinsic neuronal excitability. Thus, CS-elicited NE release in the PFC promotes a higher responsivity in the S-neurons during threat memory retrieval.

### PFC-induced activation of LC neurons during memory retrieval

Previous studies have indicated that the conditioned threat cue elevated PFC NE levels (Feenstra et al., 2001). To directly monitor changes in PFC NE level, we injected rAAV-hSyn-NE2h-WPRE-hGH virus (NE fluorescent sensors; Feng et al., 2023) into the dmPFC and monitored extracellular NE level using fiber photometry (Fig. 6A). In the conditioned mice, presentation of 30 s or 10 s CS+ led to a rapid and significant increase in the NE sensor fluorescence which decayed slowly towards pre-CS level after CS termination (Fig. 6A). In contrast, a 30 s CS- did not elicit significant change in fluorescence in the same mice (Fig. 6A), indicating that PFC NE rise is selective to CS+. In comparison, a 2 s CS+ (mimicking T-neuron activation; Fig. 3A, 3C) or 10 s CS+ led to a shorter NE increase, with the responses to 2 s CS+ longer than the CS+ duration (Fig. 6A). We observed similar response dynamics using NE sensor with medium affinity to NE (rAAV-hSyn-NE2m-WPRE-hGH; Feng et al., 2023) (Sup Fig. 6), indicating that the affinity of the NE sensor does not significantly affect the dynamic changes in NE level. A major LC input to the PFC arises from the locus coeruleus (LC). We observed a significant increase in Ca^2+^ signals to 30 s and 10 s CS+ but not to 30 s CS- in the LC-neurons expressing CaMKII-GCaMP6s virus (Fig. 6B; Florin-Lechner et al., 1996). In addition, a significant increase in Ca^2+^ signals was elicited by 2 s CS+ or 10 s CS+ (Fig. 6B). The PFC NE signals elicited by 2 s opto-stimulation of PFC-projecting LA/B neurons were similar to that by 2 s CS+ (Fig. 6C).

**Figure 6.**
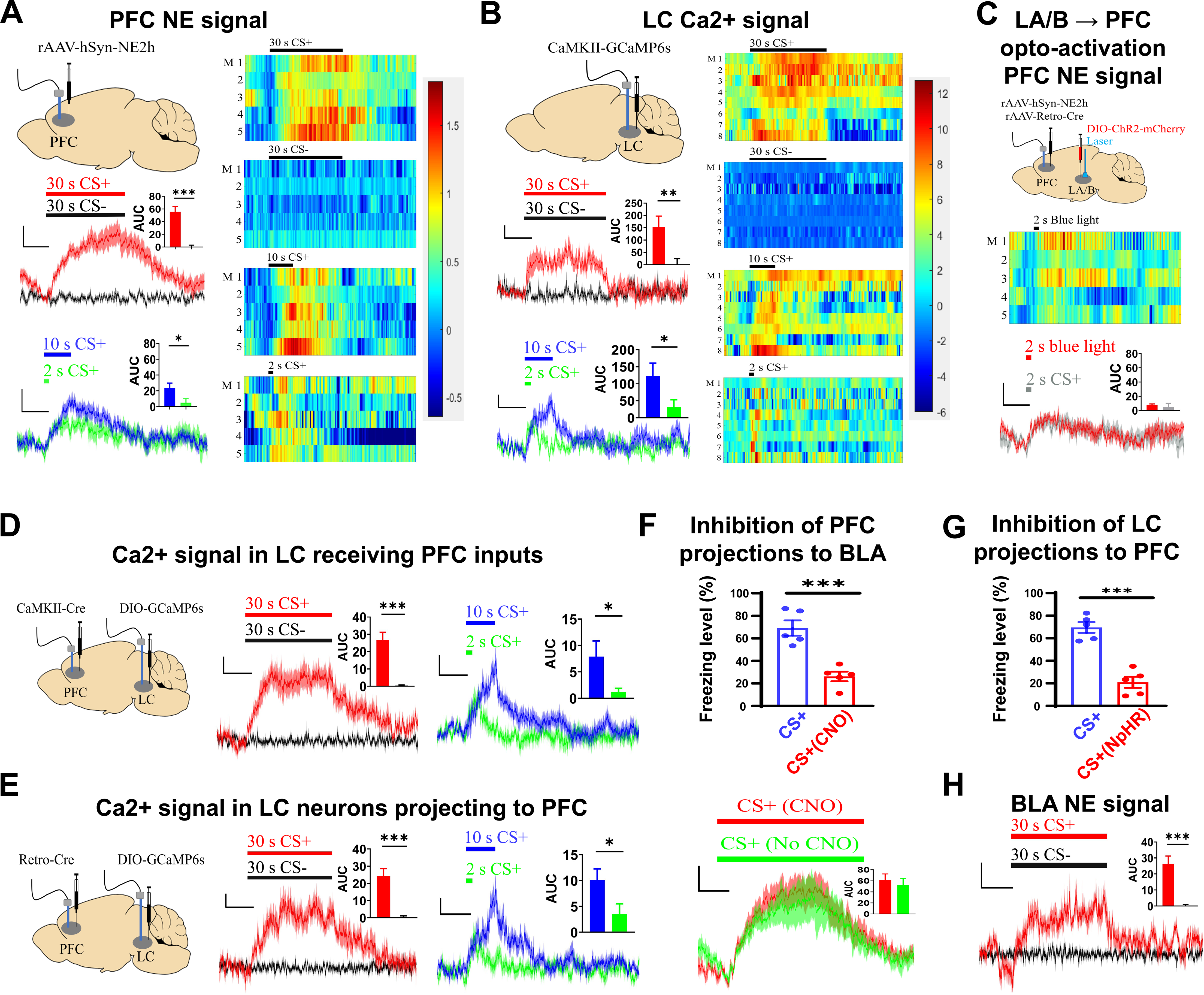
The dmPFC NE profiles associated with CS and T-neuron activation. (A) (Left) Sites of virus injections and recording (upper), and plots of NE signals with stimuli shown on the top. rAAV-hSyn-NE2h-WPRE-hGH viruses were used. Scale bars, 0.5%ΔF/F and 10 s. Inserts show the AUC of ΔF/F of PFC NE signals. (Right) Heat map showing the dmPFC NE signals to 30 s CS+, 30 s CS-, 10 s CS+ and 2 s CS+. N= 5 mice each group. (B) (Left) Sites of virus injections and recording (upper), and plots of Ca^2+^ signals with stimuli shown on top. CaMKII-GCaMP6s viruses were used. Scale bars, 2%ΔF/F and 10 s. Inserts show the AUC of ΔF/F of LC Ca^2+^ signals. (Right) Heat maps of Ca^2+^ responses in LC-neurons in response to 30 s CS+, 30 s CS-, 10 s CS+ and 2 s CS+. N= 8 mice for each group. (C) PFC NE signals to 2 s blue light stimulation of LA/B. (Upper) Sites of virus injections and recording, with rAAV-hSyn-NE2h-WPRE-hGH, rAAV-Retro-Cre and DIO-ChR2-mCherry viruses used. (Middle) Heat map. (Bottom) Traces of PFC NE responses. The 2 s CS+ response (gray, taken from Fig. 6A). N= 5 mice for each group. Scale bars, 0.5%ΔF/F and 10 s. Insert show the AUC of ΔF/F of PFC NE signals. (D) (Left) Experimental procedures for virus injections, (middle and right) plots of Ca^2+^ responses to 30 s CS+, 30 s CS-, 10 s CS+ and 2 s CS+ in LC neurons receiving PFC projections. CaMKII-Cre viruses injected in PFC and DIO-GCaMP6s viruses injected in LC. Scale bars, 0.5%ΔF/F and 10 s. N= 8 mice. Insert show the AUC of ΔF/F of Ca^2+^ signals. (E) (Left) Experimental procedures for virus injections, (middle and right) plots of Ca^2+^ responses to 30 s CS+, 30 s CS-, 10 s CS+ and 2 s CS+ in LC neurons projecting to PFC. Retro-Cre viruses injected in PFC and DIO-GCaMP6s viruses injected in LC. Scale bar, 0.5%ΔF/F and 10 s. N= 8 mice. Insert show the AUC of ΔF/F of Ca^2+^ signals. (F) (Upper) Effect of chem-inhibition of PFC-to-BLA projections on the freezing level. (Bottom) Effect of chem-inhibition of PFC-to-BLA projections on PFC NE signal. Clozapine N-oxide (CNO) was given half an hour before CS+. DIO-hM4D and NE sensor high-affinity viruses in PFC and Retro-Cre virus in BLA. Two-tailed paired t-test, CS+ vs. CS+ (CNO), P < 0.001. N= 5 mice. Insert show the AUC of ΔF/F of PFC NE signals. (G) Impact of opto-inhibiting LC-to-PFC projections on freezing level with Retro-Cre viruses injected in dmPFC and DIO-NpHR-mCherry viruses injected in LC. Comparison between freezing levels to 30 s CS+ in the presence and absence of inhibition. Two-tailed paired t-test, CS+ vs. CS+ (NpHR), P < 0.001; N= 5 mice, each group. (H) Plots of BLA NE sensor responses to 30 s CS+ and 30 s CS-. Scale bars, 0.5%ΔF/F and 10 s. N= 4 mice. Insert show the AUC of ΔF/F BLA NE responses.

The above findings indicate that activation of the PFC T-neurons may directly activate LC-neurons, which in turn project to the dmPFC and release NE (Arnsten and Goldman-Rakic, 1984; Jodo et al., 1998; Barcomb et al., 2022; Hallock et al., 2023). Supporting this, we observed prominent fluorescence in the PFC and TeA but not in BLA in mice injected with retrobeads in the LC (Sup Fig. 7). To further confirm that LC neurons that receive PFC inputs do respond to CS+, we monitored their Ca^2+^ responses to CS+ and CS- and observed selective responses to 30 s CS+ (Fig. 6D; Sup Fig. 8). In addition, these LC neurons also showed Ca^2+^ responses to 2 s and 10 s CS+, and these responses roughly matched the CS duration (Fig. 6D; Sup Fig. 8). The LC-neurons that receive PFC inputs send their projections back to dmPFC but not to LA/B (Sup Fig. 9). We observed Ca^2+^ responses to 2 s, 10 s and 30 s CS+, but not to 30 s CS, in the LC neurons that project to PFC (Fig. 6E; Sup Fig. 10), consistent with the elevated NE level to these CSs. In addition, inhibition of PFC to BLA projections significantly reduced freezing levels without affecting the PFC NE signals (Fig. 6F), consistent with these projections acting downstream of PFC-NE signaling. To further confirm the importance of the LC-PFC projections to freezing behavior, we inhibited LC-to-PFC projections by injecting retro-Cre viruses in dmPFC and DIO-NpHR- mCherry viruses in LC. We found that this inhibition significantly reduced the freezing level during CS+ retrieval (Fig. 6G), similar to that observed with propranolol, supporting the critical contribution of LC-PFC projections to freezing behavior. Since LC projections to BLA also play a prominent role in threat conditioning (Uematsu et al., 2017), we monitored the NE sensor responses in the BLA. We observed a significant and time-clocked increase in NE responses to 30 s CS+ but not to 30 s CS- in the BLA, with faster rise and decay compared to NE signals in the PFC (Fig. 6H).

### A rise in PFC NE level opens a short window for transition to behavior during memory retrieval

The T-neuron responses last ∼ 2 s, but the S-neuron responses and freezing roughly match the duration of CS+. This brief spiking pattern suggests that T-neuron activity may gate or enable the ensuing responses. This raises the question of whether CS following T-neuron activation can sustain freezing. We addressed this question using a hybrid CS signal: a brief (2 s) CS+ followed immediately by a CS- (28 s) (Fig. 7A). The 2 s CS+ mimics the duration of T-neuron responses and limits the S-neuron responses (Fig. 3D). Interestingly, freezing levels (Fig. 7A), T-neuron and S-neuron responses (Fig. 7B), were all indistinguishable between the hybrid CS+/CS- and a 30 s CS+, in the same set of mice. In the above experiments, CS+ may activate other neurons besides T-neurons. To address this possibility, we replaced CS+ with a 2 s opto-activation of T-neurons (L/CS- hybrid, Fig. 7C). We found that freezing levels (Fig. 7C), responses in T-neurons and S-neurons (Fig. 7D), were all indistinguishable between L/CS- and CS+. Note that the frequency and pattern used for CS- in these experiments were very different from those used for CS+ (see Methods), and CS- alone did not elicit significant freezing. Control experiments using light or 2 s CS+ only were not conducted since 2 sec is too short to measure freezing levels accurately (see Methods). The freezing level elicited by 2 s opto-stimulation of PFC neurons that receive LA/B followed by 28 s CS- was indistinguishable from that elicited by 30 s CS+ (Fig. 7E). These results strongly indicate that a brief CS+ presentation or LA/B activity is sufficient to elicit significant S-neuron responses and freezing when followed any CS (sound). The S-neuron responses/freezing match the duration of the CS used (i.e., CS+/CS- hybrid), supporting the critical role of sensory/sound inputs.

**Figure 7.**
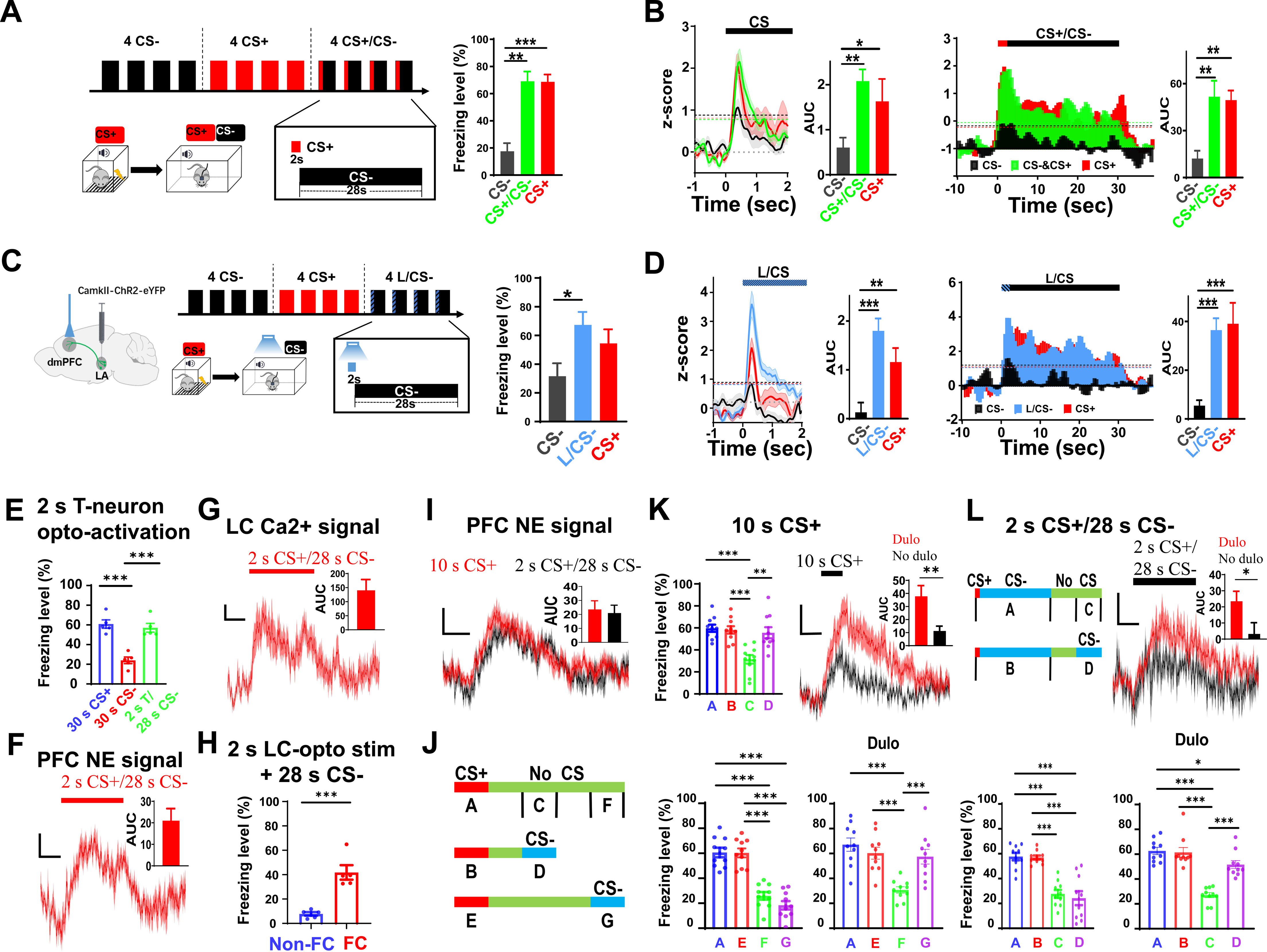
Brief activation of dmPFC T-neurons opens a window for transition to freezing behavior. (A) (Left) Experimental protocol and stimulus patterns used and (right) corresponding freezing levels. One-way RM ANOVA, F (2,8) = 0.163, Bonferroni’s posttest; CS- vs. CS+/CS-, P < 0.01; CS- vs. CS+, P < 0.001; N= 10 mice. (B) (Left) Spike rates in T-neurons during CS retrieval (One-way RM ANOVA, F (2, 72) = 4.699, Bonferroni’s posttest; CS- vs. CS+/CS-, P < 0.01; CS- vs. CS+, P < 0.05; n= 25 units/10 mice). (Right) Spike rates in S-neurons during CS retrieval (One-way RM ANOVA, F (2, 33) = 2.092, Bonferroni’s posttest; CS- vs. CS+/CS-, P < 0.01; CS- vs. CS+, P < 0.01; n= 12 units/10 mice). Inserts show the AUC (z-score) of CS- elicited responses with three CSs. (C) (Left) Experimental protocol for opto-activation of LA/B-dmPFC connections for 2 s followed by CS- (at CS onset) during CS retrieval, and (right) corresponding freezing levels. Mice were conditioned with CS+ only. One-way RM ANOVA, F (2,18) = 3.807, Bonferroni’s posttest; T-neurons, CS- vs. L/CS-, P < 0.05; N= 8 mice. (D) (Left) Spike rates in T-neurons (One-way RM ANOVA, F (2, 42) = 6.67, Bonferroni’s posttest; CS- vs. L/CS-, P < 0.001; CS- vs. CS+, P < 0.05; n= 15 units/6 mice). (Right) Spike rates in S-neurons (One-way RM ANOVA, F (2, 18) = 5.099, Bonferroni’s posttest; CS- vs. L/CS-, P < 0.001; CS- vs. CS+, P < 0.001; n= 7 units/6 mice). (E) Freezing to 30 s CS+, 30 s CS- and 2 s opto-activation of T-neurons/28 s CS-. Two-tailed paired t-test, 30 s CS+ vs. 30 s CS-, P < 0.001; 30 s CS- vs. 2 s T-opto/28 s CS-, P < 0.001. N= 5 mice. (F) NE signals in dmPFC elicited by 2 s CS+/28 s CS-. Scale bars, 0.5%ΔF/F and 10s. N= 5 mice. Insert show the AUC of ΔF/F of NE signals. (G) LC Ca^2+^ signals elicited by 2 s CS+/28 s CS-. Scale bars, 2%ΔF/F and 10 s. N= 8 mice. Insert show the AUC of ΔF/F of Ca^2+^ signals. (H) Freezing to 2 s opto-activation of LC neurons/28 s CS- in unconditioned/conditioned mice. Two-tailed paired t-test, Non-FC vs. FC, P < 0.001. N= 5 mice. (I) 10 s CS+ and 2 s CS+/28 s CS- elicited PFC NE signal. Insert show the AUC of ΔF/F of NE signals. (J) Experimental procedure for testing the impact of CS- on freezing level when given after of break after 10 s CS+ termination (CS+/B/CS-). Different sections during the experiments have been named A, B….G, to mark the period for the analysis of freezing. (K) (Upper left and lower left) Freezing levels elicited by the protocol. Two-tailed paired t-test, Ctrl (0 - 10 s) vs. Ctrl (20 - 30 s), P < 0.001; CS+/CS- (0 - 10 s) vs. Ctrl (20 - 30 s), P < 0.001; Ctrl (20 - 30 s) vs. CS+/CS- (20 - 30 s), P < 0.01; Ctrl (0 - 10 s) vs. Ctrl (40 - 50 s), P < 0.001; Ctrl (0 - 10 s) vs. CS+/CS- (40 - 50 s), P < 0.001; CS+/CS- (0 - 10 s) vs. Ctrl (40 - 50 s), P < 0.001; CS+/CS- (0 - 10 s) vs. CS+/CS- (40 - 50 s), P < 0.001. N= 10 mice. (Upper right) PFC NE signal during 10 s CS+ enhanced in the presence of NE uptake inhibitor duloxetine (10 mg/kg, i.p.). (Lower right) Freezing levels elicited by CS+/CS- (40 - 50 s in the presence of duloxetine (10 mg/kg, i.p.). Two-tailed paired t-test, Ctrl (0 - 10 s) vs. Ctrl (40 - 50 s), P < 0.001; CS+/CS- (0 - 10 s) vs. Ctrl (40 - 50 s), P < 0.001; Ctrl (40 - 50 s) vs. CS+/CS- (40 - 50 s), P < 0.001. Scale bars, 0.5%ΔF/F and 10 s. N= 10 mice. Insert show the AUC of ΔF/F of PFC NE signals. (L) (Upper left) Experimental procedure for testing the impact of CS- on freezing level when given after a break after 2 s CS+/28 s CS- termination. (Lower left) Freezing levels elicited by the protocol. Two-tailed paired t-test, Ctrl (0 - 30 s) vs. Ctrl (40 - 50 s), P < 0.001; Ctrl (0 - 30 s) vs. CS+/CS- (40 - 50 s), P < 0.001; CS+/CS- (0 - 30 s) vs. Ctrl (40 - 50 s), P < 0.001; CS+/CS- (0 - 30 s) vs. CS+/CS- (40 - 50 s), P < 0.001. N= 10 mice. (Upper right) PFC NE signals during 2 s CS+/28 s CS- in the presence of duloxetine (10 mg/kg, i.p.). (Lower right) Freezing levels elicited by the protocol in the presence of duloxetine (10 mg/kg, i.p.). Two-tailed paired t-test, Ctrl (0 - 30 s) vs. Ctrl (40 - 50 s), P < 0.001; Ctrl (0 - 30 s) vs. CS+/CS- (40 - 50 s), P < 0.05; CS+/CS- (0 - 30 s) vs. Ctrl (40 - 50 s), P < 0.001; Ctrl (40 - 50 s) vs. CS+/CS-(40 - 50 s), P < 0.001. Scale bars, 0.5%ΔF/F and 10 s. N= 10 mice. Insert show the AUC of ΔF/F of PFC NE signals.

Since significant LC-NE signaling is observed with 2 s CS+ or opto-activation of T-neurons, and activation of β-ARs is required to sustain S-neuron response and freezing, hybrid CS+/CS- may enable S-neuron responses and freezing by eliciting significant PFC NE rise. Supporting this hypothesis, the PFC NE signals elicited by the 2 s CS+/28 s CS- remained high during the entire CS presentation (Fig. 7F; Sup Fig. 11). Consistently, sustained LC Ca^2+^ signals were also observed in response to 2 s CS+/28 s CS- (Fig. 7G; Sup Fig. 11). In addition, Ca^2+^ responses to 2 s CS+/28 s CS-were also observed in LC-neurons receiving PFC inputs, or projecting to PFC (Sup Fig. 12). Importantly, 2 s opto-stimulation of LC neurons followed by 28 s CS- was sufficient to elicit significant freezing (Fig. 7H). Significant BLA NE responses were also elicited by 2 s CS+/28 s CS- or 30 s CS+ (Sup Fig. 13). Since CS- alone does not elicit a significant increase in PFC NE, this finding further suggests that brief CS+ inputs enable LC neurons to respond to CS- to maintain NE release in PFC. Consistently, we observed robust responses in LC neurons to opto-stimulation of TeA inputs in slices (Sup Fig. 14).

Is the PFC NE level both necessary and sufficient for eliciting freezing in conditioned mice? We noticed that PFC NE signals for 10 s CS+ and 2 s CS+/28 s CS- were similar in amplitude and dynamics (Fig. 7I), and 2 s CS+/28 s CS- elicited freezing indistinguishable from that of 30 s CS+. If a high enough PFC NE level is sufficient to elicit freezing, we reasoned that a CS- given between 20 and 30 s in the 10 s CS+ experiments (section D in Fig. 7J) may elicit significant freezing. We observed significant freezing with CS- given between 20 and 30 s but not between 40 and 50 s after CS+ onset (section G in Fig. 7J) (Fig. 7K). The absence of freezing during section C (Fig. 7J, 7K) was likely due to the absence of CS in that experiment. We further tested the relationship between PFC NE level and freezing by altering PFC NE level using NE reuptake inhibitor duloxetine. We observed higher peak and longer decay in the PFC NE signal for 10 s CS+ (Fig. 7K), but no significant responses to 30 s CS- in mice injected with duloxetine (Sup Fig. 15). In the presence of duloxetine, significant freezing occurred between 40 and 50 s after CS+ onset (section G in Fig. 7J) (Fig. 7K). When we repeated the above experiments using 2 s CS+/28 s CS-, no significant freezing was observed when a CS- was given between 40 and 50 s after CS+ onset (section G) (Fig. 7L), while significant freezing was observed with the same CS- when mice were injected with duloxetine (Fig. 7L). These results collectively indicate that freezing may readily be elicited when a PFC NE level is sufficiently high and a CS is present.

## Discussion

In this study, we have demonstrated that three essential components act in concert to enable the successful transition from a formed conditioned memory to observable behaviors. The dmPFC T-neurons function as memory units and a hub of integration to determine whether a memory is expressed. The dmPFC S-neurons are activated by dmPFC T-neurons and TeA (sensory) inputs, and they convert transient PFC responses to sustained ones to match cue duration. Elevated dmPFC NE level is elicited by threat cue (CS+), and it is required to enhance S-neuron responses and the occurrence of freezing behavior. This highly interactive process enables a memory to fit the specific CSs and this transition process is subjected to modulation by the emotional and bodily states present during memory retrieval (a model is presented in Sup Fig. 16 and 17, respectively).

For a memory to best guide behavior, it should (1) encode the key information extracted from past/learned experience and (2) specify the elicited behavior to best fit the context/condition during memory retrieval. In Pavlovian threat conditioning, the CS predicts the occurrence of US (Pavlov, 1927; Rescorla, 1988; Pearce and Bouton, 2001; Fanselow et al., 2015). This CS-US association, being the gist of a threat memory, is encoded by the LA/B neurons. The transition from memory to behavior requires: (1) activated memory, which ensures the specificity for the elicited behavior (i.e., by CS+ but not CS-); (2) sensory (TeA) inputs, which specifies the duration of the responses; (3) signals representing bodily/emotional states for fine-tuning responses according to the context/conditions. Effective integration of the above three factors assures the specificity, accuracy, and relevance of the memory-triggered behavior.

### T-neurons as memory units

During retrieval of threat memory, CS+ elicits a transient response in LA/B neurons and dmPFC T-neurons. The specificity of T-neuron responses to conditioned cues is reflected in their positive correlation to freezing levels and the absence of their responses to safety or neutral cues (Yan et al., 2019; 2021; Fig. 2). Inhibition of T- neuron activation reduces/abolishes S-neuron activation and freezing, supporting that their activation is required for the transition to behavior. These T-neurons receive powerful inhibition from local PV-neurons, which receive extensive inputs from various brain regions such as the ventral hippocampus, ventral tegmental area, and paraventricular thalamus (Sotres-Bayon et al., 2012; Courtin et al., 2014; Yan et al., 2019; Wang et al., 2023). These brain regions represent or respond to safety cues, cues with low CS-US association or stress (Yan et al., 2019; 2021; Wang et al., 2023; Wang et al., in press). The T-neurons display similar responses to both CS+ and CS- during a state of generalized freezing (Yan et al., 2019), consistent with dmPFC contributing to fear generalization (Lissek, Bradford et al., 2014; Vieira et al., 2015). The T-neurons encode threat memory using a population-based mechanism, with each neuron responding in an all-or-none manner.

### S-neurons as behavior units

Activation of S-neurons is required for transitioning from the transient T-neuron responses to sustained freezing (Fig. 3). S-neurons integrate a few key factors and fill in the critically missing information (such as sensory information). S-neurons use an activity-based encoding mechanism for this transition. The magnitude of the S-neuron responses is determined by T-neuron activity, NE signals, and the intrinsic excitability of S-neurons. In contrast, the duration of the S-neuron responses is determined by the TeA inputs. Opto-activation of T-neurons resulted in a few distinct steps in the S- neuron responses, suggesting S-neurons integrate T-neuron responses. Although both T- and S-neurons are present in dmPFC and likely close to each other spatially, S- neurons differ from T-neurons in a few important ways: (1) inputs: S-neurons receive inputs from the TeA and T-neurons, while T-neurons receive inputs from LA/B and local GABAergic neurons. (2) Outputs: S-neurons project to BLA while T-neurons project to S-neurons and LC. (3) Electrophysiological properties: S-neurons have higher input resistance, longer spike half-width, lower spike amplitude, and conditioning-dependent synaptic modifications. (4) Genes: T- and S-neurons show differential expression levels in Ca^2+^ channels, cell-adhesion molecules and β-ARs. These differences support their distinct contributions to the transition from memory to behavior.

### LC-NE system as a critical modulator

NE signaling is the third critical player in transitioning from memory to behavior. Current literature supports the view that the LC-NE system is important for the formation, consolidation, and extinction of conditioned threat responses (Uematsu et al., 2017; Giustino and Maren, 2018; Likhtik and Johansen, 2019). However, the exact contribution of how LC/NE enhances the transition from memory to behavior is poorly understood. There are many unanswered questions, including (1) what inputs to the LC neurons trigger the NE release; (2) the selectivity of LC neuron activation in regard to the conditioned cues; (3) whether LC/NE exerts the classic amplification effect seen in the sensory system; (4) whether NE modifies the retrieved memory.

Our findings demonstrate the LC contribution to the memory-elicited behavior. First, activation of PFC-projecting LC neurons enables or even enhances the execution of behavior (freezing) via enhanced synaptic and neuronal responses that result in enhanced excitation leading to behavior. This finding extends the proposed LC’s adaptive gain function to include the solicitation of the behavior by memories (Aston-Jones and Cohen, 2005). These activated LC neurons function as a filter to select the sensory inputs with behavioral importance, and in our study, CS+. More importantly, our findings suggest that LC neurons gate such behavior because their activation allows the ensuing behaviorally unimportant cues to maintain the elicited behavior. Second, LC neurons execute the above function in a modular manner. The participating LC neurons are activated by the PFC T-neurons during the threat memory retrieval, and they further project to PFC. Third, the enhanced LC function is mediated by increased LC neuron activation and net enhancement of PFC neuron responses mediated by β-ARs. Importantly, rapid plasticity in the LC neurons is likely engaged during this process. The plasticity is evidenced by the observation that CS- elicits strong responses in LC neurons when it follows a brief CS+ but not by itself.

Activation of LC neurons amplifies the S-neuron responses in a similar manner to those shown for the sensory inputs in previous studies (Aston-Jones et al., 1997; 1999; Bouret and Sara, 2004; 2005). Activation of dmPFC β-ARs is required to boost dmPFC S-neuron activity and freezing level, consistent with previous studies in mice (Burgos-Robles et al., 2009; Pendyam et al., 2013) and humans (Kroes et al., 2016). These findings are also consistent with the general notion that LC phasic activation and increased PFC NE level facilitate the appropriate behavioral responses to imperative stimuli (Aston-Jones and Cohen, 2005; Nieuwenhuis et al., 2010), and NE signaling in the central amygdala modulates conditioned threat responses (Gu et al., 2020). This action of β-ARs is consistent with their expression pattern in the mPFC (Rainbow et al., 1984; Booze et al., 1993; Nicholas et al., 1993). Our finding of β- ARs enhancing the postsynaptic neuron activity via enhanced excitatory inputs, reduced inhibitory inputs, and enhanced intrinsic excitability, is consistent with previous reports and implicating that NE increases signal-to-noise ratio (Ferry et al., 1999; Polack et al., 2013; Schiemann et al., 2015; Skelly et al., 2017; Bacon et al., 2020). Although our NE fluorescence signals cannot be readily converted into NE concentrations, it has been shown that the extracellular NE concentrations in the cerebral cortex are linearly related to LC output across a range of tonic LC activity (Berridge and Abercrombie, 1999; Florin-Lechner et al., 1996). It should be noted that β-ARs do not significantly impact T-neurons’ responses or their properties. Therefore, the role of β-ARs appears limited to facilitating the transition step by mostly affecting S-neurons, consistent with previous findings (Sara, 2009). The Glutamate Amplifies Noradrenergic Effects (GANE) model proposes that local glutamate release in the vicinity of LC neuron terminals enhances the effects of NE released from these terminals (Mather et al., 2016). This is an interesting and important mechanism to be tested in future studies.

Activation of threat memory activates the LC-neurons via PFC-to-LC projections, consistent with previous studies and our retrograde tracing results (Arnsten and Goldman-Rakic, 1984; Jodo et al., 1998; Barcomb et al., 2022; Hallock et al., 2023). The selective increase in CS+-elicited PFC NE level is similar to the enhanced responses of LC-NE neurons to aversive stimuli after threat conditioning (Uematsu et al., 2017), which may have an evolutionary advantage by enabling rapid responses to potential threats (Hughes et al., 2007). This mode of action aligns with the gain increase/control model of LC function (Aston-Jones and Cohen, 2005), with rapidly enhanced LC neuronal activity amplifying the responses of the target neurons, such as S-neurons. Our results indicate that a certain level of PFC NE needs to be reached for freezing to occur, although no quantitative relationship between the two parameters can be deduced using the current method. It is essential to notice that a sensory cue (CS) is required to trigger S-neuron responses and freezing. This observation suggests that activation of LC neurons and the resulted high NE level puts the brain into an alert/vigilant state during which the relevant sensory cues readily elicit behavior. Here, the relevant sensory cues are not restricted to what is encountered during conditioning/learning, but instead can be generalized, such as to the same sensory modality. This notion of a temporary vigilance/alertness state is consistent with the LC/NE associated with vigilance and anxiety (Poe et al., 2020; Berrocoso et al., 2022; Bouras et al., 2023). We must emphasize that the neuromodulators released during this time window mostly target the memory-elicited behavior rather than the memory itself. This also sets the therapeutic limits for targeting such neuromodulators.

The observation that CS- following a brief CS+ or PFC T-activation can elicit responses in LC neurons while unable to do so on its own suggests a rapid plasticity in the LC neurons, likely associated with the activation of their PFC inputs. This plasticity of LC-neurons is similar to that previous demonstration of rapid plasticity induced by sensory stimulation in the adult LC neurons (Martins and Froemke, 2015) and stress-induced long-lasting enhancement in the excitability of developing LC- neurons (Borodovitsyna et al., 2018). The nature of this LC neuron plasticity is worthy of further investigation. The reduced discrimination between CS+ and CS- (comparing LC neuron responses between 30 s CS+, 30 s CS- and 2 s CS+/28 s CS-) is reminiscent of previous findings wherein a generalized increase in LC/NE system elevates network gain without discrimination between inputs (Usher et al., 1999; Aston-Jones and Cohen, 2005), and hence rendering the targeted circuits more responsive to a wide range of stimuli. There are two points worthy of notice: (1) the LC responses and PFC NE elevation to CS+ are substantially higher than that to CS-; (2) the less-discriminative enhancement in LC neuron responses requires the gating by selective inputs (CS+). This finding suggests an interesting possibility of retrieved memory to open a window during which emotion-like responses may be present.

The outputs of LC-NE neurons are modular. For example, LC neuron functions differ significantly between those projecting to PFC and BLA (Chandler at al., 2014; Schwarz et al., 2015; Hirschberg et al., 2017; Uematsu et al., 2017; Chandler at al., 2019; Plummer et al., 2020; Poe et al., 2020). The LC projections to auditory regions may explain the observed elevation in dmPFC NE level during CS- presentation. Activation of LC-neurons is known to enhance sensory signaling via either synaptic plasticity (Martins and Froemke, 2015) or postsynaptic effect on ARs (Aston-Jones and Waterhouse, 2016; Vincent Breton-Provencher et al., 2021). Other potential targets of LC, such as the amygdala or bed nucleus of the stria terminalis, are known to mediate anxiety or stress (Davis, 1992; Walker et al., 2009; Roozendaal et al., 2009; Adhikari, 2014). It remains to be tested whether the significant elevation in BLA NE signals we observed during memory retravel is associated with higher vigilance or stress (Goldstein et al., 1996).

Taken together, the roles of LC/NE in the transition from memory to behavior are two-fold: it enhances the responses to behaviorally important cues, which is consistent with the gain control theory (Aston-Jones and Cohen, 2005), and it engages an emotion-like process that readies the brain to respond rapidly once the relevant cues appear. This latter function is consistent with LC’s activation when a task involves emotional cues (Sterpenich et al., 2006), and it is worthy of further exploration.

## Summary

Our findings suggest that the transition from memory to behavior is an elaborate process requiring integration of processes/factors presenting memory, sensory inputs (CS), and the internal state (NE and likely other modulators) for best fitting to the CS encountered and the bodily state at the time of memory retrieval. This integration provides another layer of modulation, allowing further fine-tuning at the transition stage from memory to behavior.

## Acknowledgments

This work is supported by grants from Shenzhen-Hong Kong Institute of Brain Science-Shenzhen Fundamental Research Institutions (2023SHIBS0004). We thank the members of the Zhou lab for the helpful discussion.

## Author contributions

Supervision, Q.Z; conceptualization, Q.Z, T.W, Y.L; methodology, Q.Z, T.W, X.Z, H.D, D.X, R.Y, Y.Z, T.L and Y.L; project administration, Q.Z; experiment and analysis, T.W, X.Z, H.D, T.L, D.X and Q.Z; writing – original draft, Q.Z, T.W; writing – review & editing, Q.Z, T.W, X.Z, H.D. Y.L, and W-J,G.; funding acquisition, Q.Z and Y.L.

## Declaration of interests

All authors declare no competing interests.

## Supplemental figures

**S-Figure 1.**
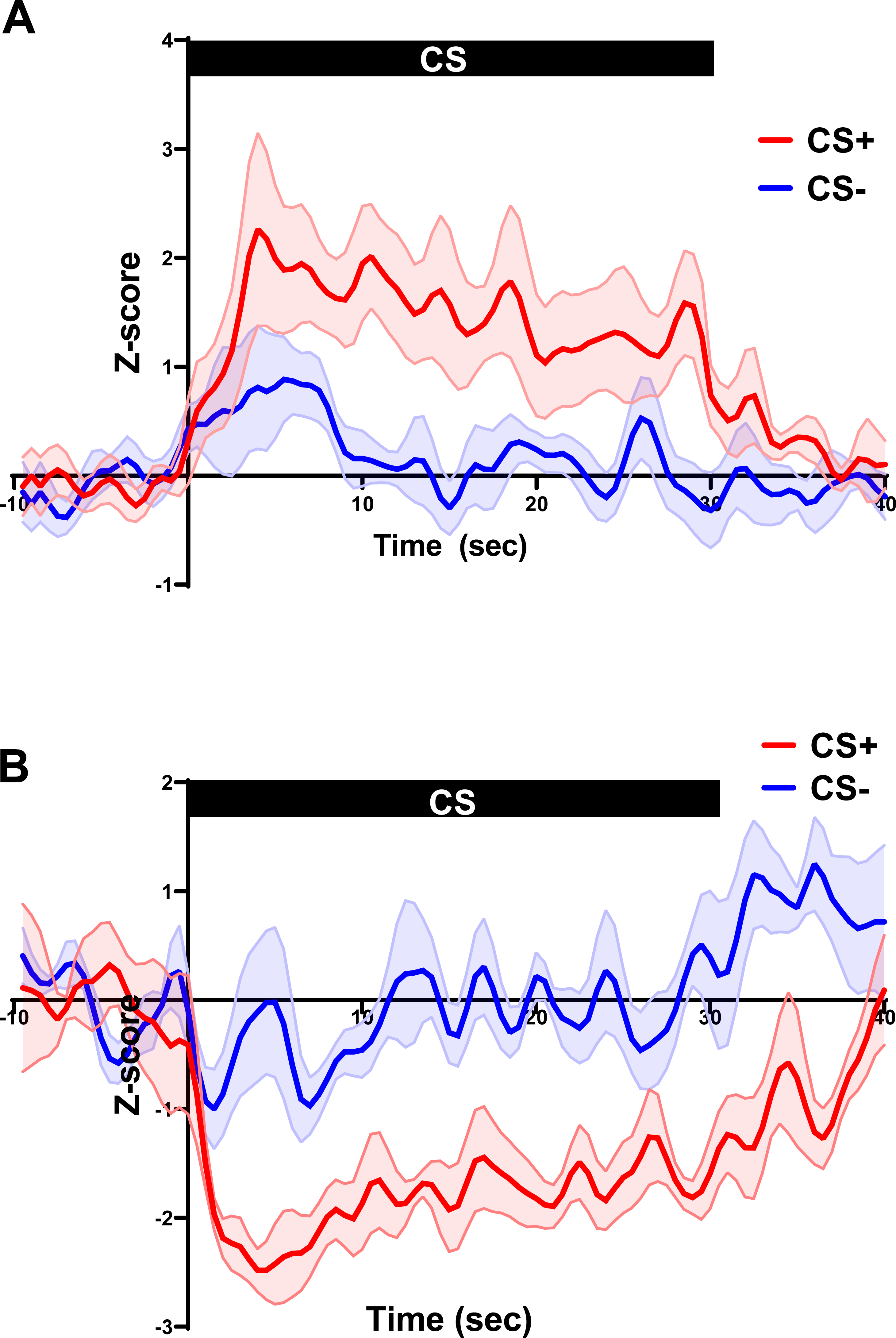
Responses in S-neurons to CS+ and CS- during retrieval. (A) Selectively increased sustained responses to CS+ and no response to CS- in the S- neurons. n= 9 units/5 mice. (B) Selectively decreased sustained responses to CS+ and no response to CS- in the S- neurons. n= 5 units/5 mice.

**S-Figure 2.**
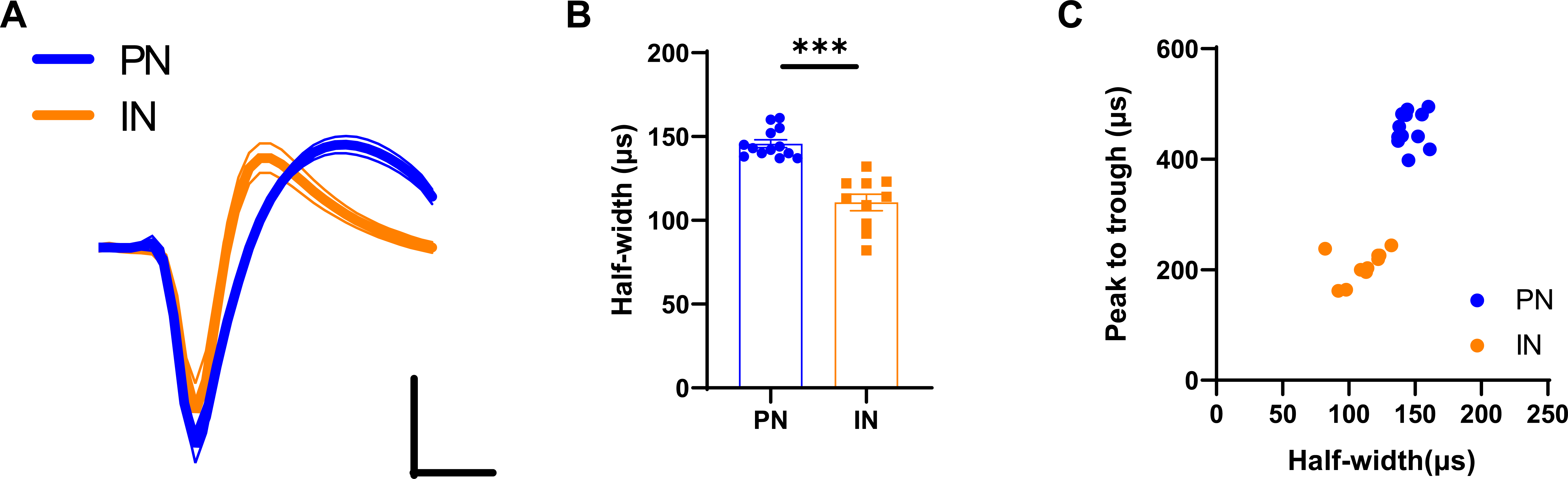
Distinguishing inhibitory and excitatory neurons based on their spike waveforms. (A) The averaged voltage waveforms of all INs and PNs. Thick lines represent the average while thin lines the SEM. Scale bars, 50 µV and 250 µs. (B) Quantification of half-width of INs and PNs. PNs, 13 units/5 mice; Ins, 10 units/6 mice; two-tailed Student’s t-test, *, P < 0.001. (C) Plot for peak to trough and half-width of all units.

**S-Figure 3.**
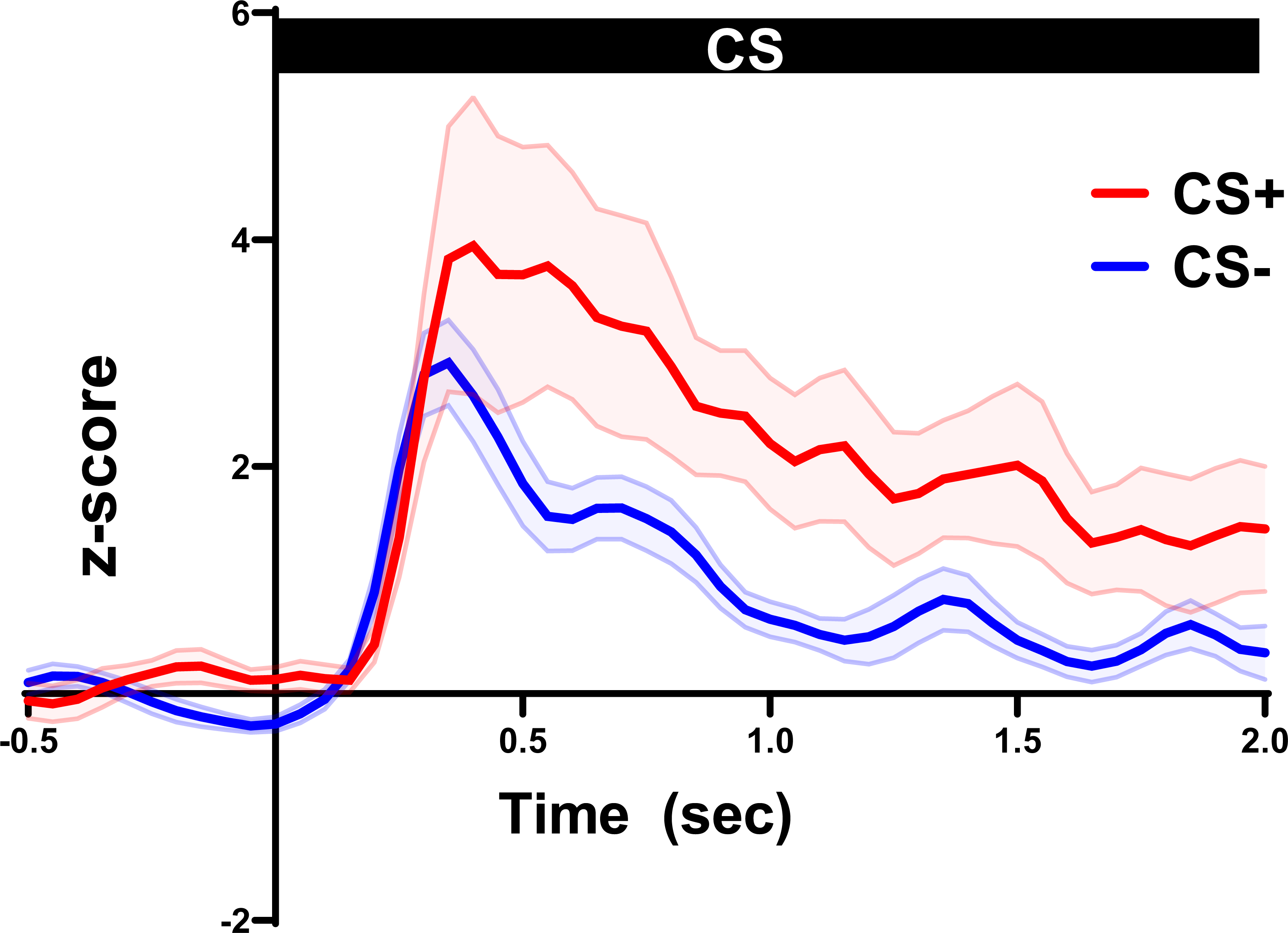
Transient responses to CS+ and CS- in the LA/B neurons. n=23 units/8 mice

**S-Figure 4.**
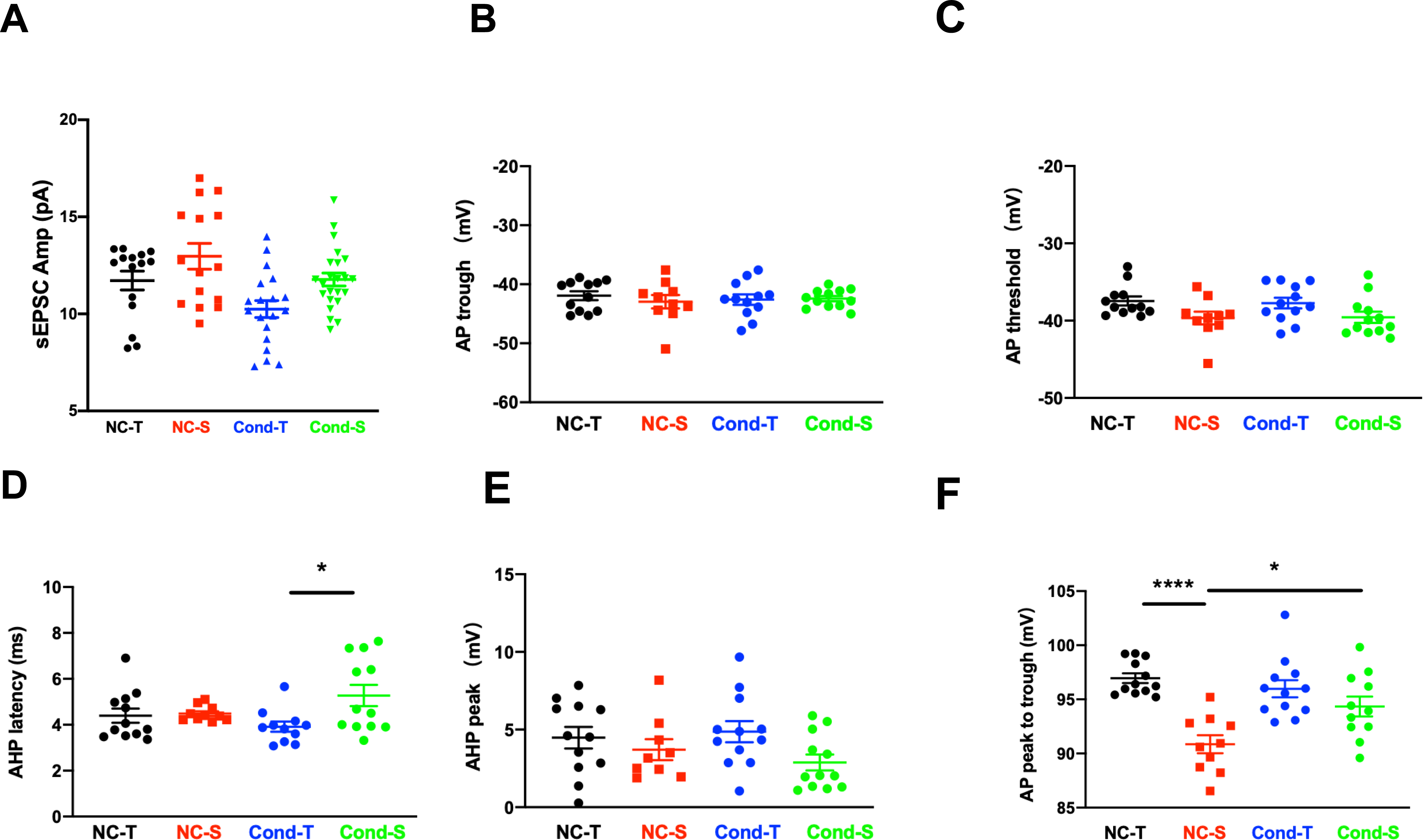
Electrophysiological properties of T-neurons and S-neurons recorded in PFC slices. (A) sEPSC amplitude in dmPFC T-neurons and S-neurons. One-way RM ANOVA, F (2,53) = 7.804, Bonferroni’s posttest, P > 0.05; n= 15 cells/4 mice (NC-T), 15 cells/4 mice (NC-S), 20 cells/4 mice (Cond-T), 24 cells/6 mice (Cond-S). (B) Spike trough in T- and S-neurons. One-way RM ANOVA, F (3, 46) = 2.734, P < 0.05, Bonferroni’s posttest; NC-S vs. Cond-S, P < 0.05; 12 cells/4 mice (NC-T), 10 cells/4 mice (NC-S), 12 cells/4 mice (Cond-T), 12 cells/4 mice (Cond-S). (C) Spike threshold in T- and S-neurons. One-way RM ANOVA, F (3, 54) = 9.415, P < 0.001, Bonferroni’s posttest; NC-S vs. Cond-S, P < 0.001; 12 cells/4 mice (NC-T), 10 cells/4 mice (NC-S), 12 cells/4 mice (Cond-T), 12 cells/4 mice (Cond-S). (D) AHP latency in T- and S-neurons. One-way RM ANOVA, F (3, 41) = 3.199, P < 0.05, Bonferroni’s posttest; NC-S vs. Cond-S, P < 0.05; 12 cells/4 mice (NC-T), 10 cells/4 mice (NC-S), 12 cells/4 mice (Cond-T), 12 cells/4 mice (Cond-S). (E) AHP peak in T- and S-neurons. One-way RM ANOVA, F (3, 41) = 1.969, P > 0.05, Bonferroni’s posttest; 12 cells/4 mice (NC-T), 10 cells/4 mice (NC-S), 12 cells/4 mice (Cond-T), 12 cells/6 mice (Cond-S). (F) Spike threshold in T- and S-neurons. One-way RM ANOVA, F (3, 41) = 11.84, P < 0.001, Bonferroni’s posttest; NC-S vs. Cond-S, P < 0.001; NC-S vs. Cond-S, P < 0.05; 12 cells/4 mice (NC-T), 10 cells/4 mice (NC-S), 12 cells/4 mice (Cond-T), 11 cells/4 mice (Cond-S).

**S-Figure 5.**
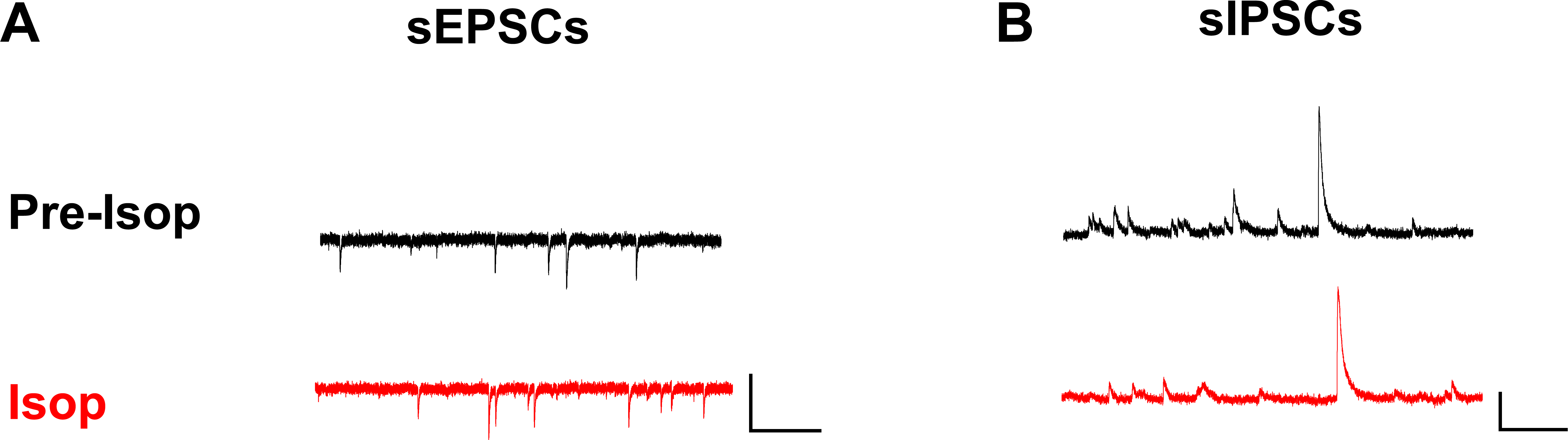
Impact of Isop on T-neurons. (A) Sample recordings of sEPSCs in the T-neurons before and after bath application of Isop. Scale bars, 50 pA and 1 s. (B) Sample recordings of sIPSCs in the T-neurons before and after bath application of Isop. Scale bars, 50 pA and 500 ms.

**S-Figure 6.**
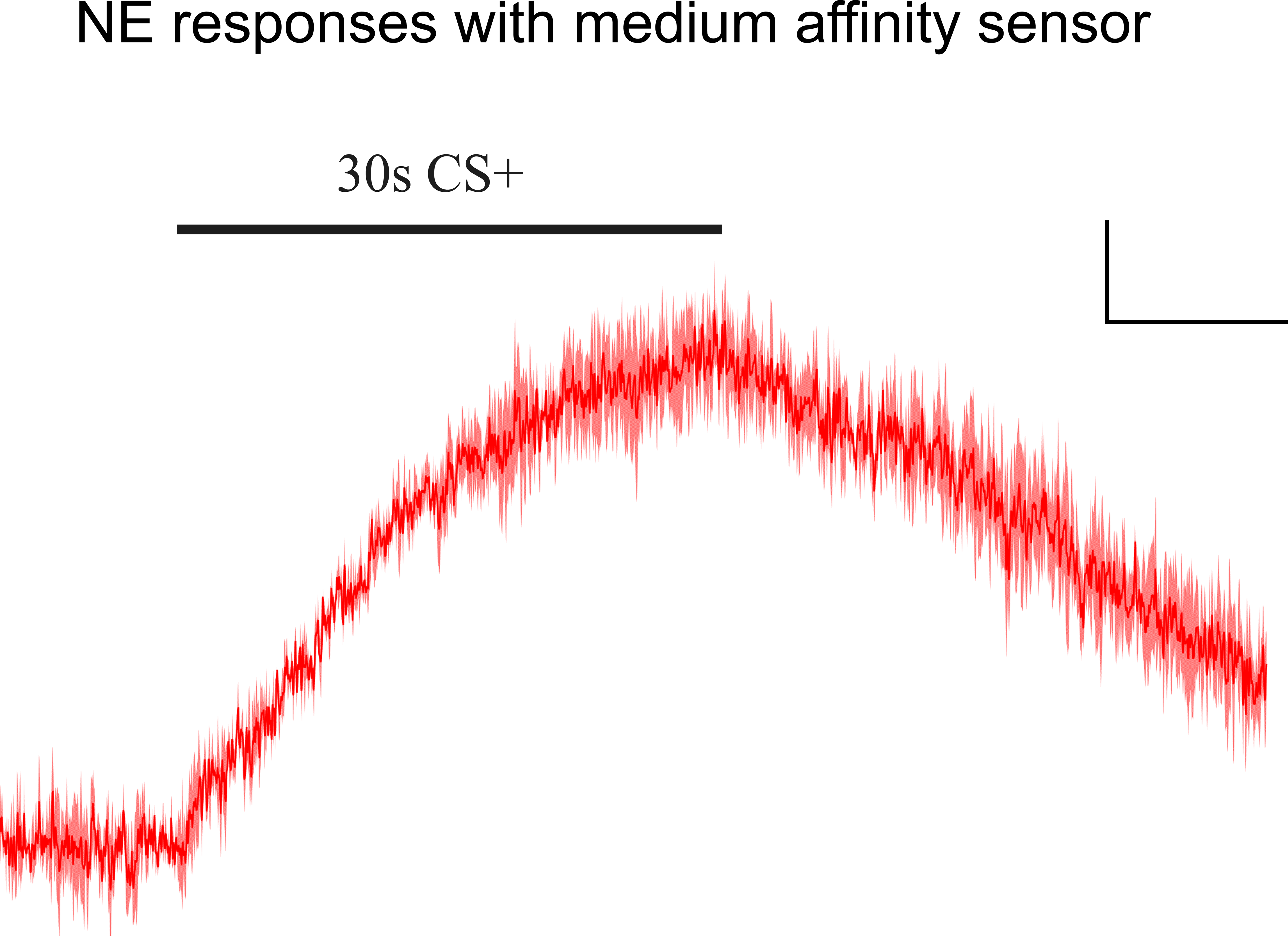
NE responses to 30 s CS+ detected using medium affinity (encoded by rAAV-hSyn-NE2m-WPRE-hGH virus) in PFC. Scale bars, 0.5%ΔF/F and 10 s. Trace is an average of 3 trials.

**S-Figure 7.**
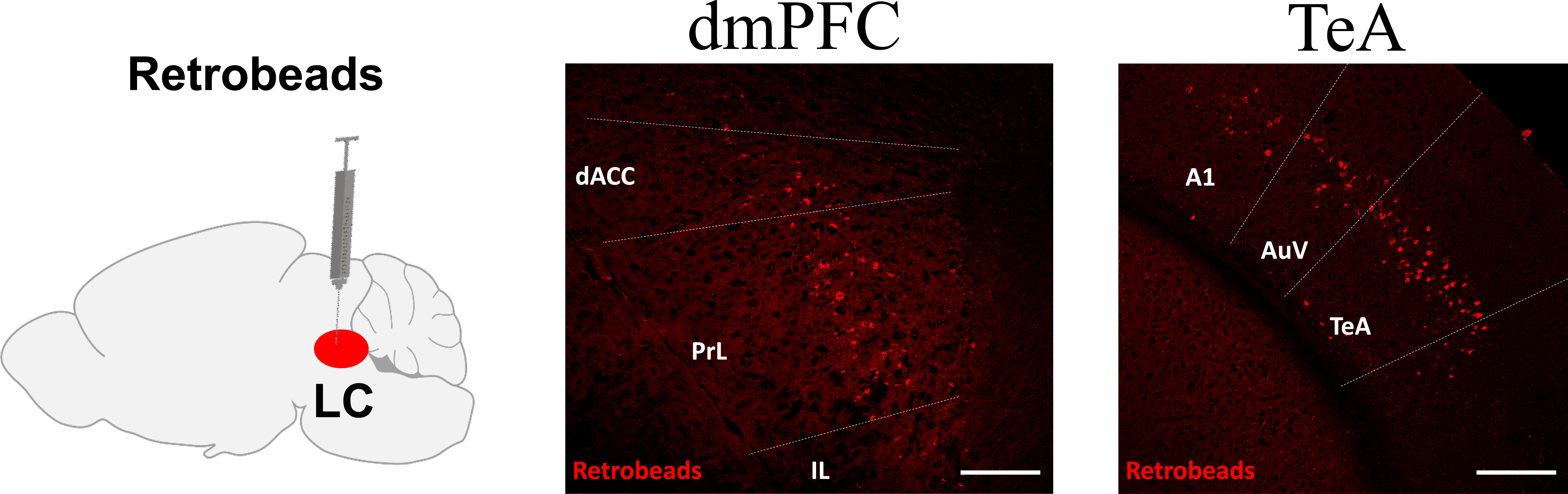
Major projections to LC neurons identified using retrobeads injected in the LC. Scale bar, 200 μm.

**S-Figure 8.**
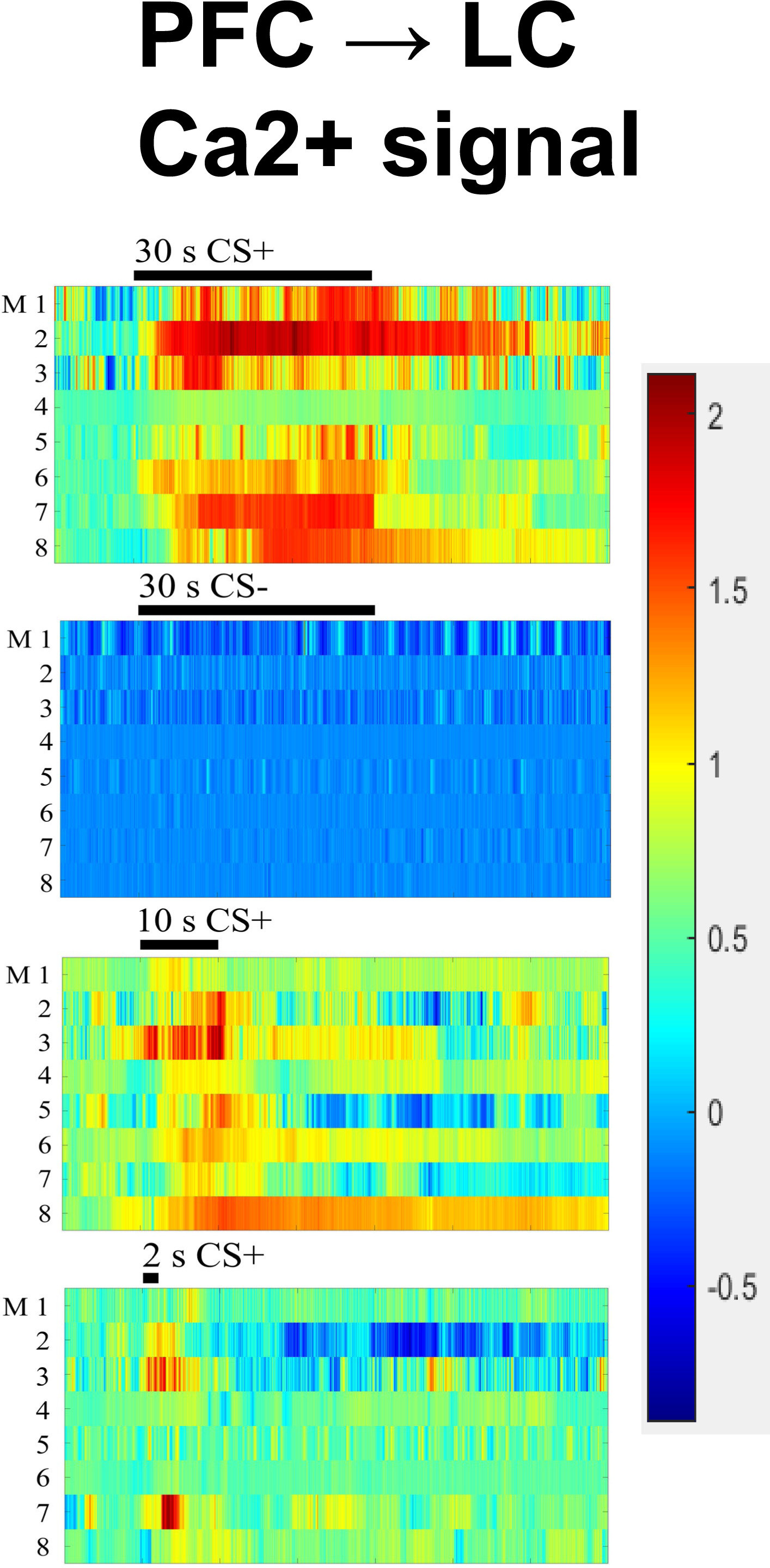
Heat map of Ca^2+^ responses to 4 sets of CSs in LC neurons receiving PFC projections. CaMKII-Cre viruses injected in PFC and DIO-GCaMP6s viruses injected in LC. N= 8 mice.

**S-Figure 9.**
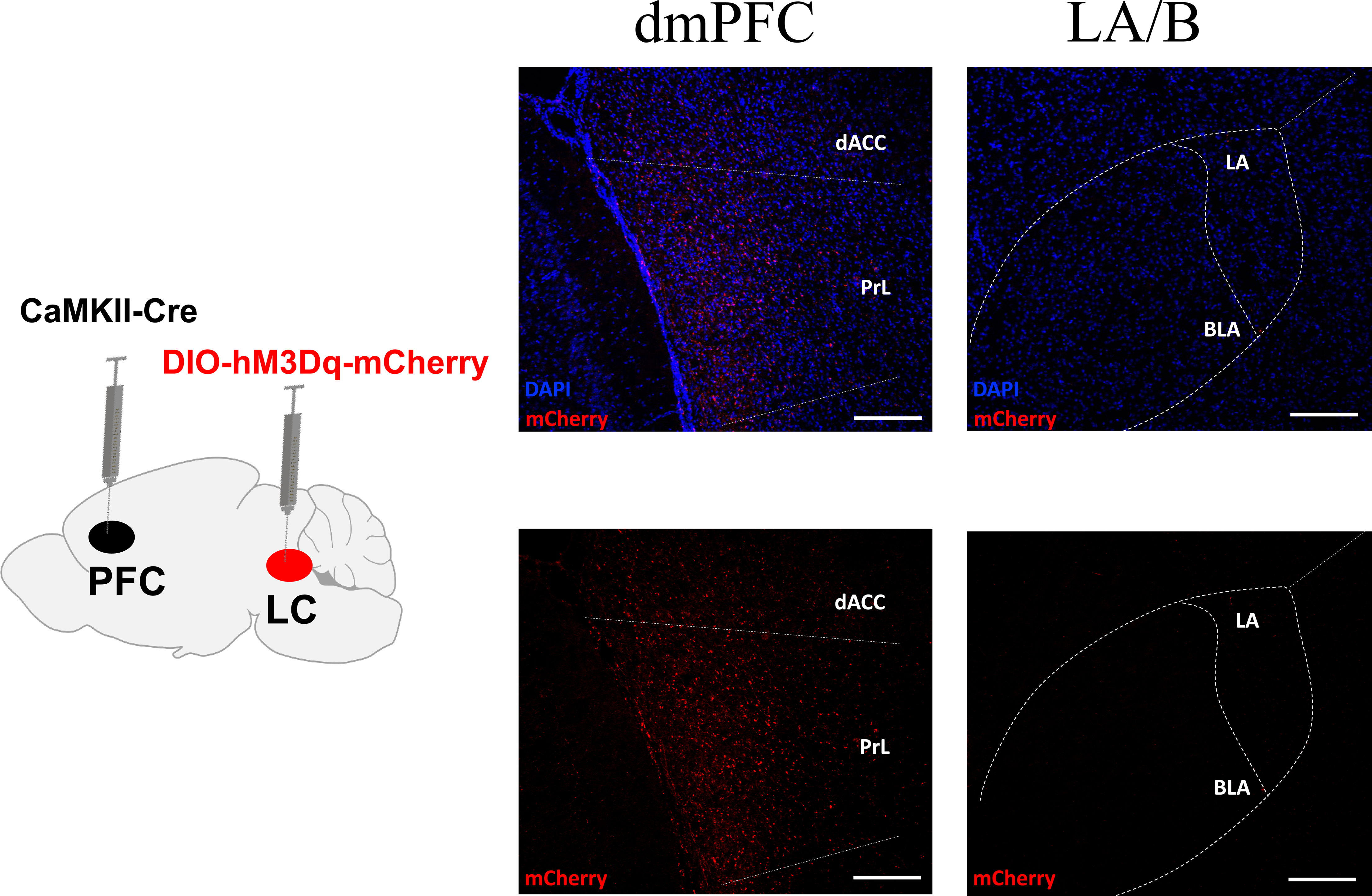
(Left) Experimental paradigm for examining outputs from LC neurons that receive PFC projections, using injection of CaMKII-Cre virus in dmPFC and DIO-hM3Dq-mCherry virus in LC. (Right) Presence of the mCherry fluorescence in dmPFC and not in LA/B. Scale bars, 200 μm.

**S-Figure 10.**
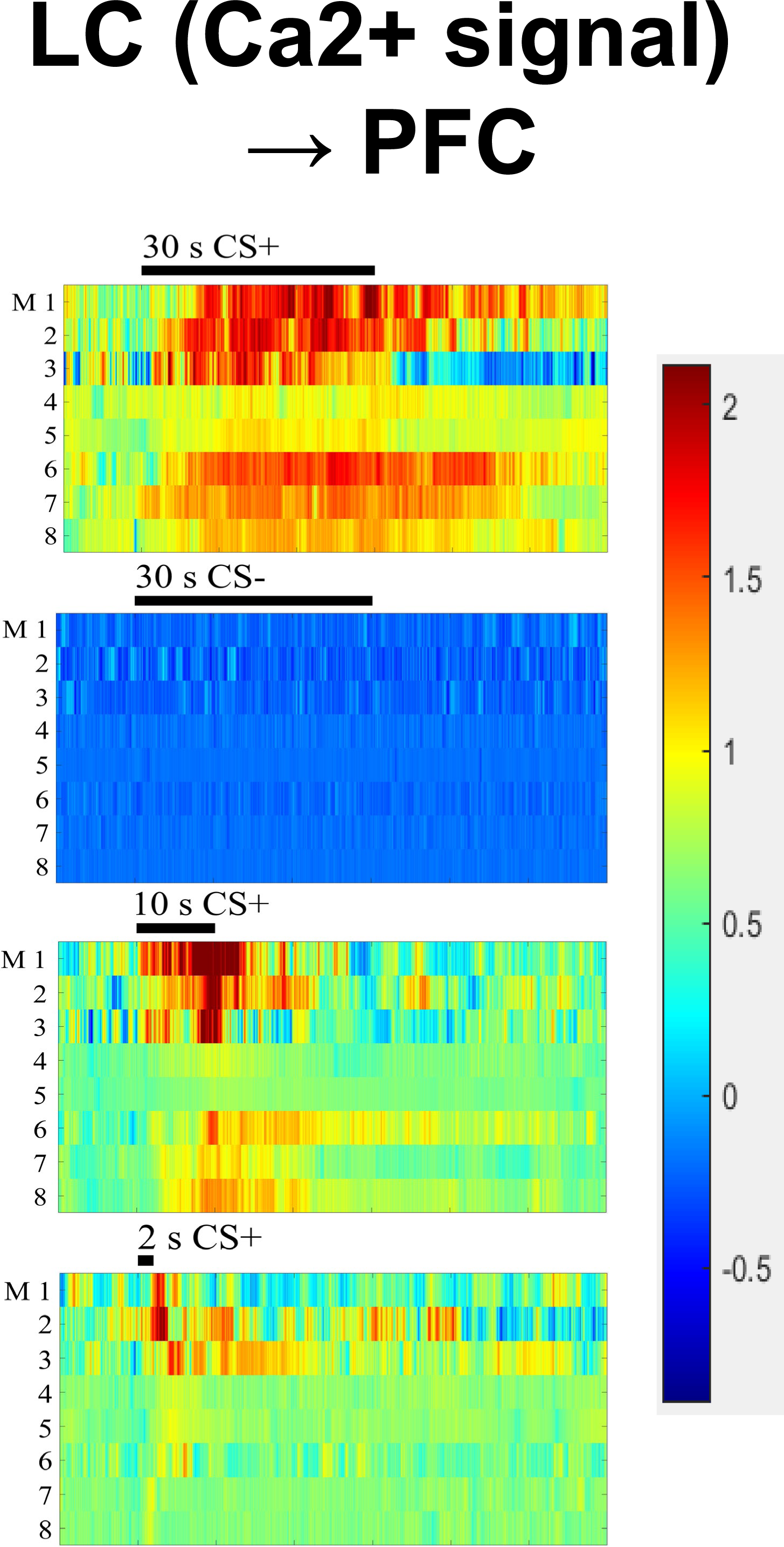
Heat map of Ca^2+^ responses to 4 sets of CSs in LC neurons projecting to PFC. Retro-Cre viruses injected in PFC and DIO-GCaMP6s viruses injected in LC. N= 8 mice.

**S-Figure 11.**
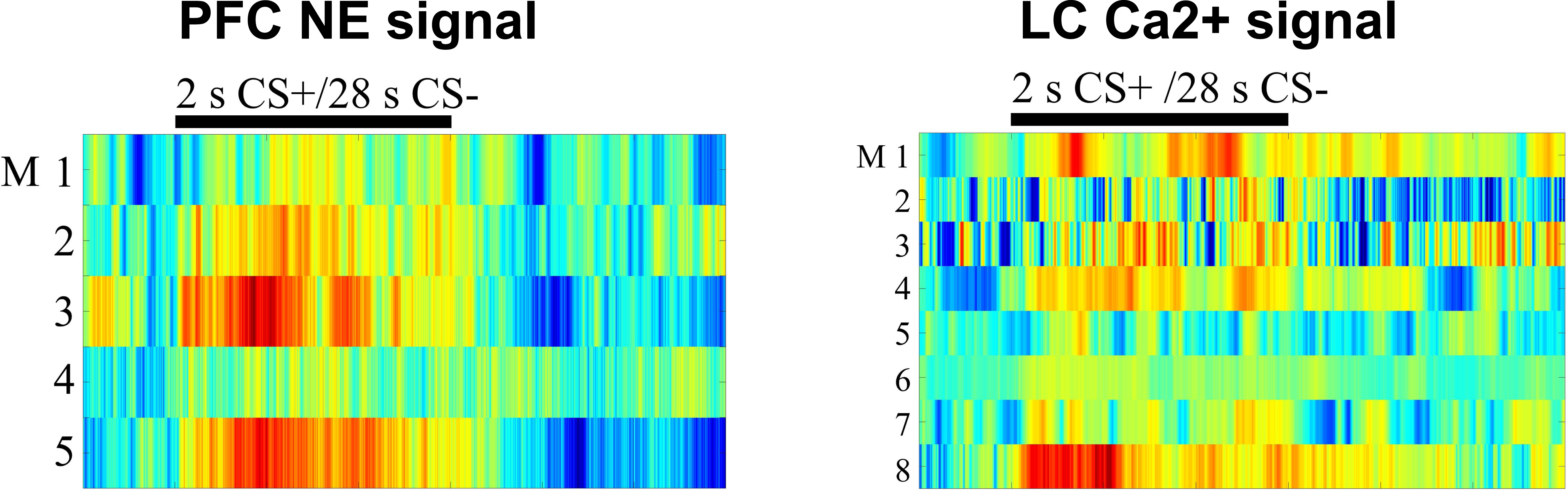
(Left) Heat map of PFC NE responses to 2 s CS+/28 s CS-. (Right) Heat map of Ca^2+^ responses to 2 s CS+/28 s CS-.

**S-Figure 12.**
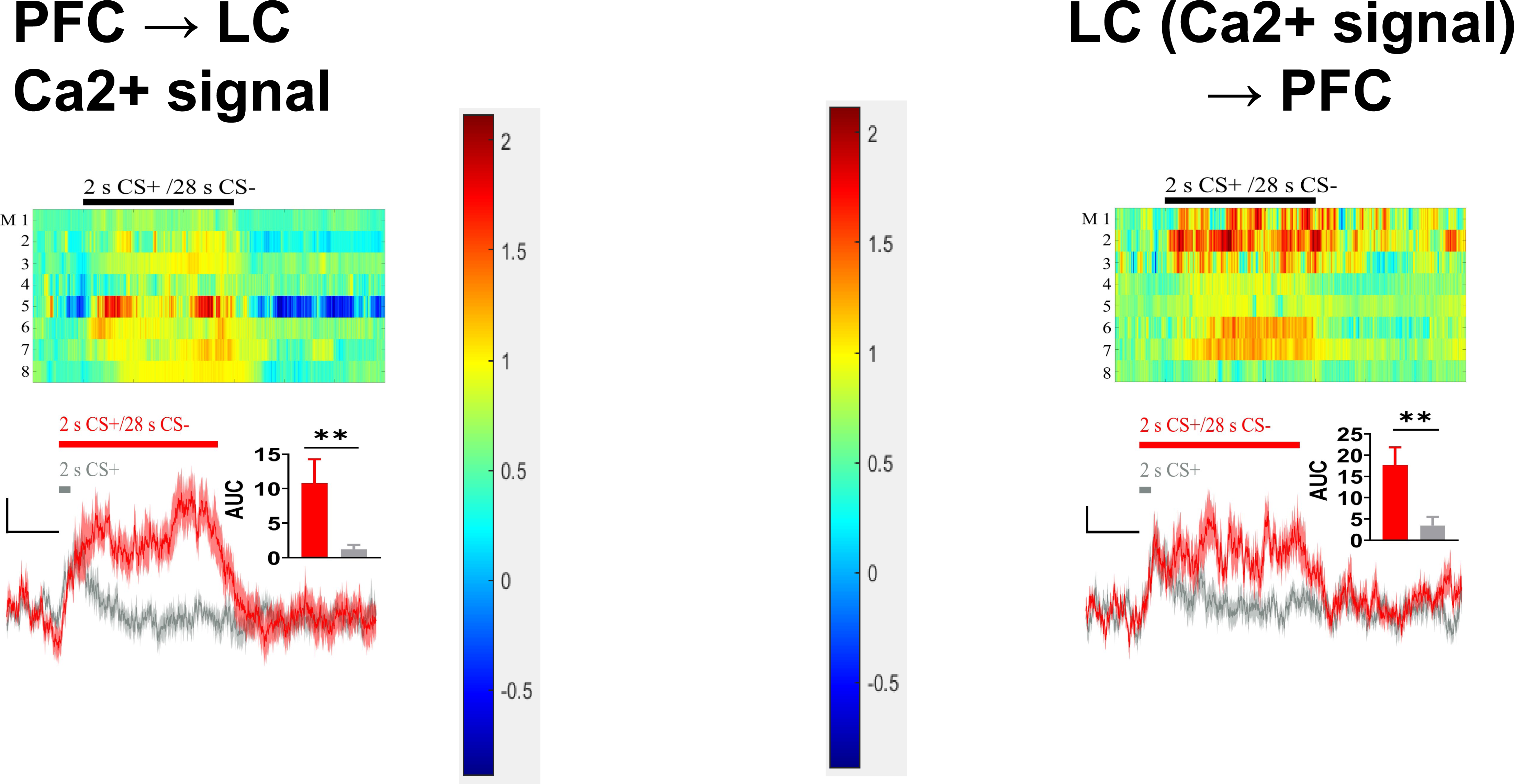
Heat map and plot of Ca^2+^ signals elicited by 2 s CS+/28 s CS- and 2 s CS+ in LC neurons receiving PFC projections (left) and projecting to PFC (right). N = 8 mice.

**S-Figure 13.**
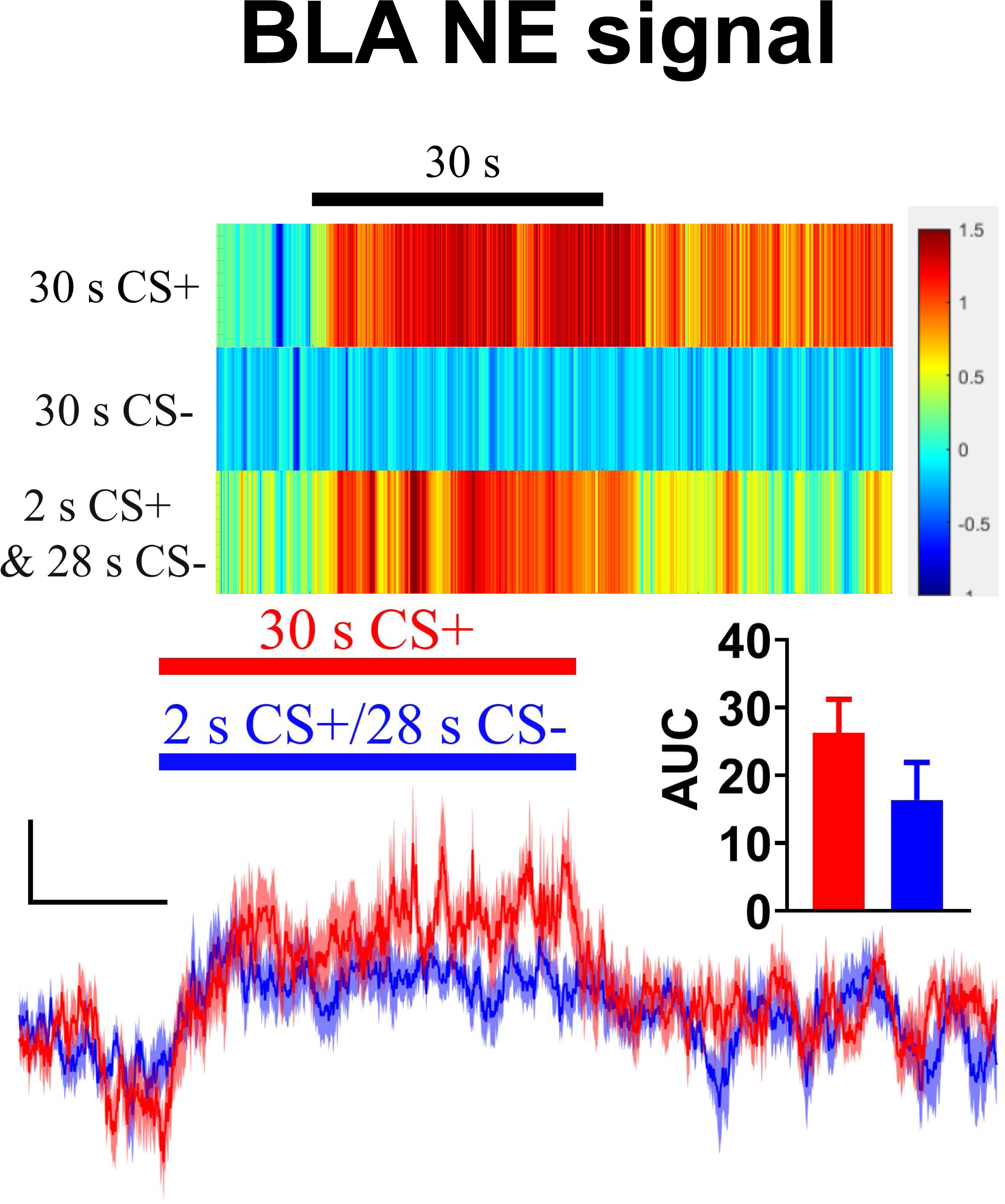
(Upper) Representative heat map of BLA NE sensor responses to 30 s CS+, 30 s CS- and 2 s CS+/28 s CS-, and (bottom) plots of BLA NE signals to 30 s CS+ and 2 s CS+/28 s CS-. Scale bars, 0.5%ΔF/F and 10 s. N= 4 mice. Insert show the AUC of ΔF/F of BLA NE signals.

**S-Figure 14.**
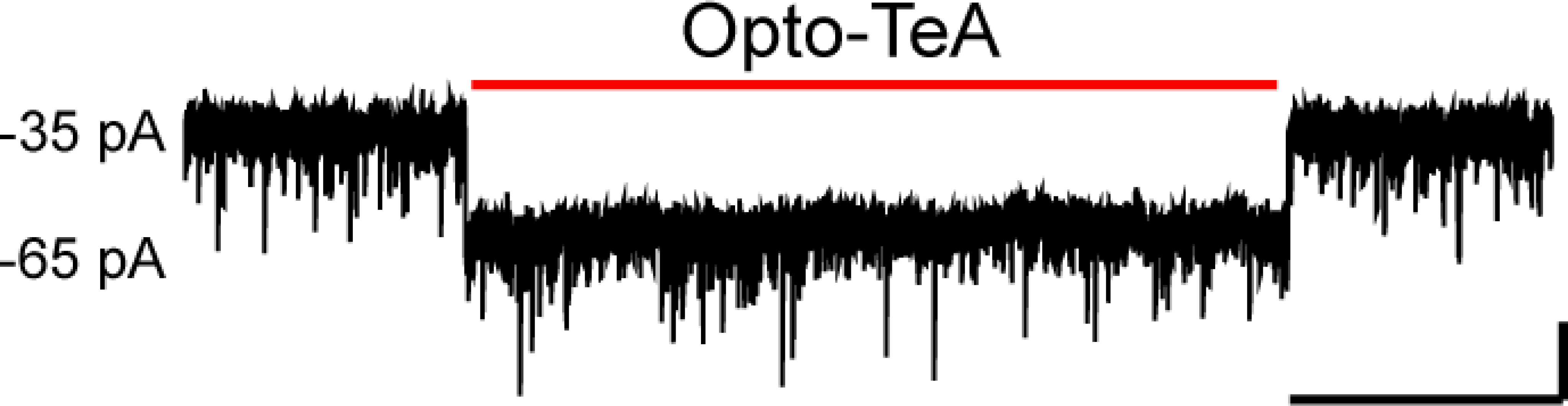
Responses of LC neurons to opto-stimulation of TeA inputs in LC slice. Sample recording of LC neurons held in voltage clamp (at ™70 mV), with opto-stimulation marked by red line. Scale bars, 20 pA and 5 s.

**S-Figure 15.**
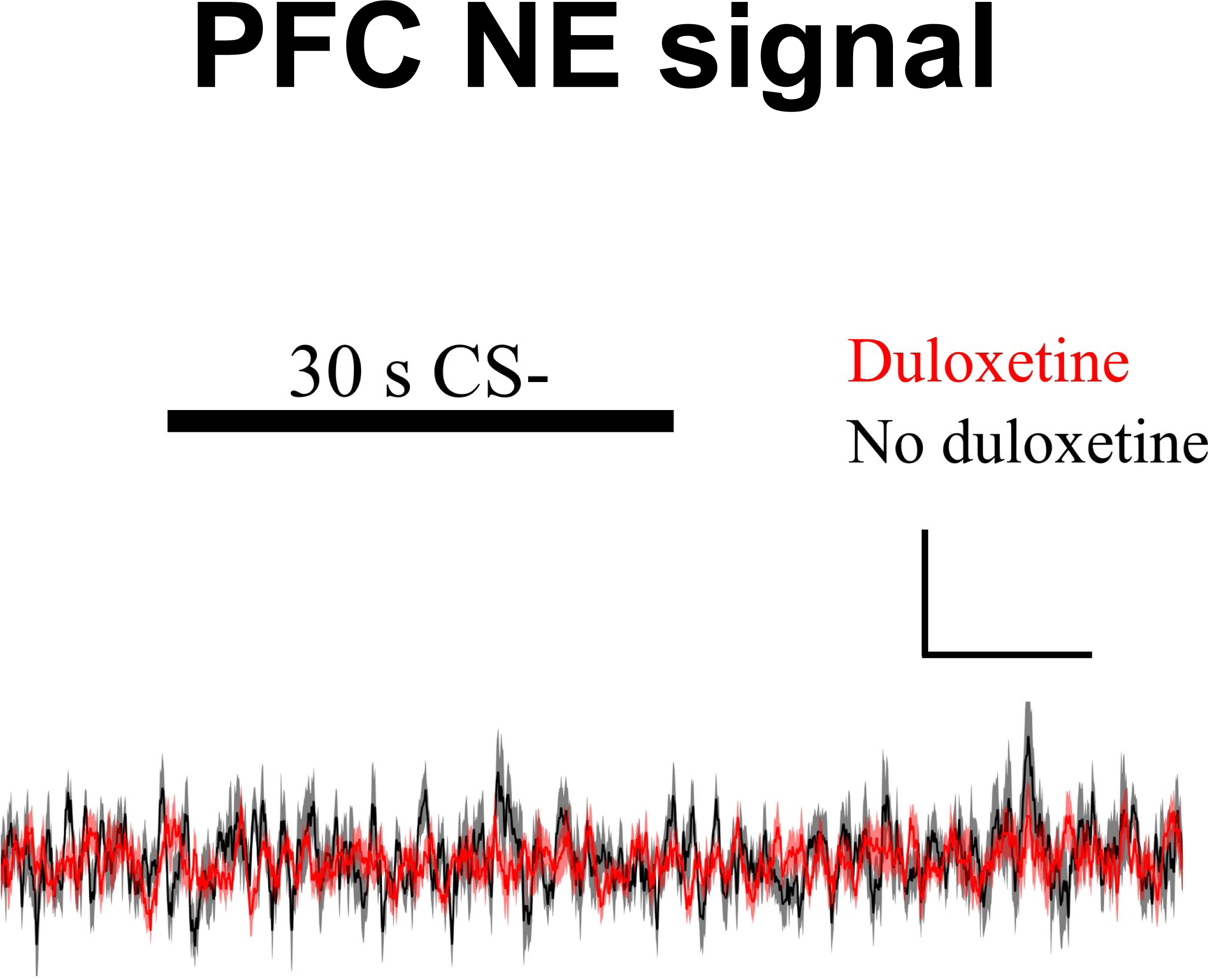
PFC NE signal during 30 s CS- in the presence of NE uptake inhibitor duloxetine (10 mg/kg, i.p.). Scale bars, 0.5%ΔF/F and 10 s. N= 10 mice.

**S-Figure 16.**
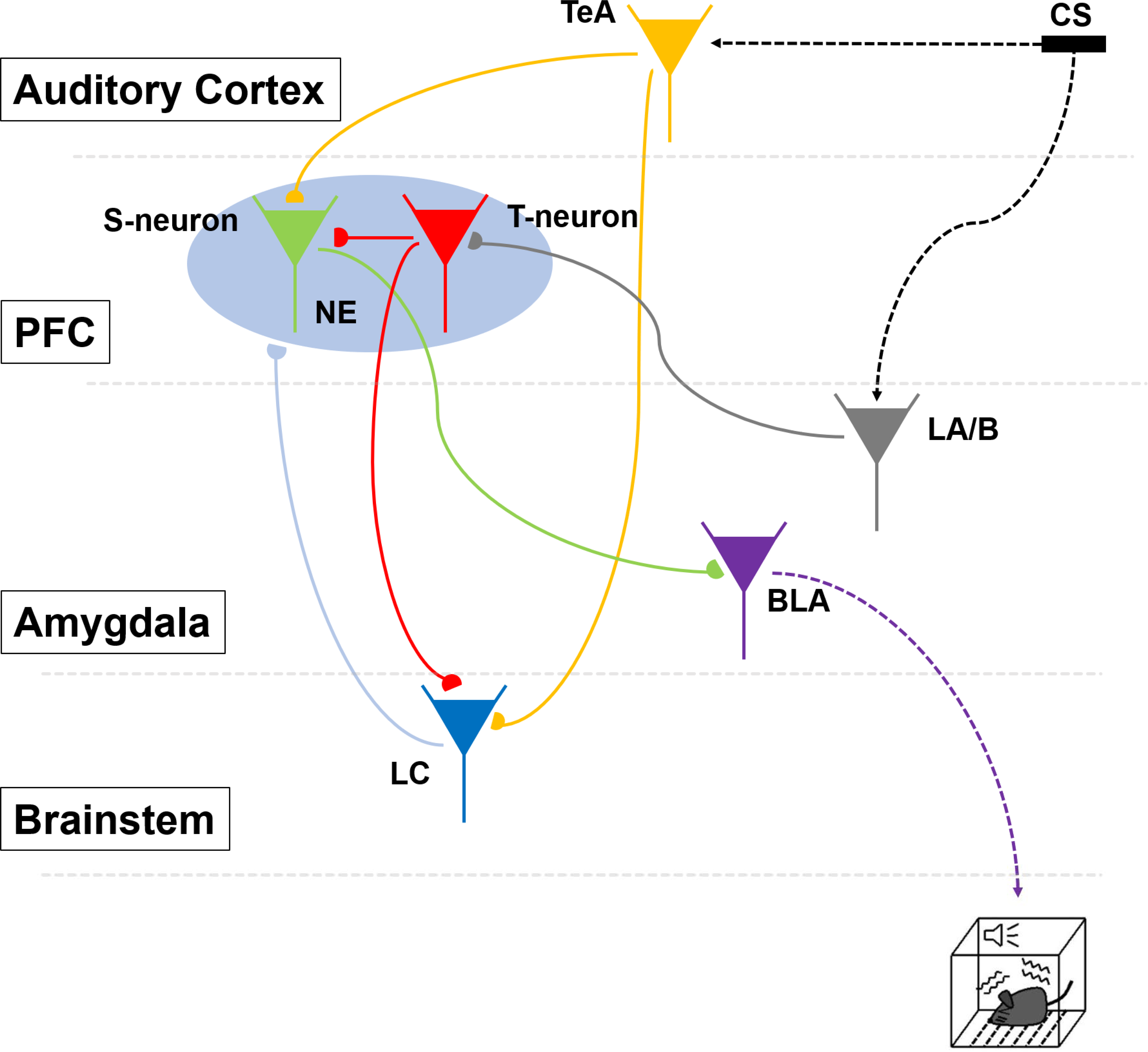
Model of the neural circuit exhibiting the information flows and modulation from conditioned memory to freezing behavior. Presentation of CS+ activates LA/B neurons, which, in turn, activates PFC T-neurons. T-neurons activate PFC S-neurons and LC neurons. S-neurons receive inputs from the TeA and send outputs to the BLA to enable freezing behavior. Activated LC neurons project to PFC and release NE.

**S-Figure 17.**
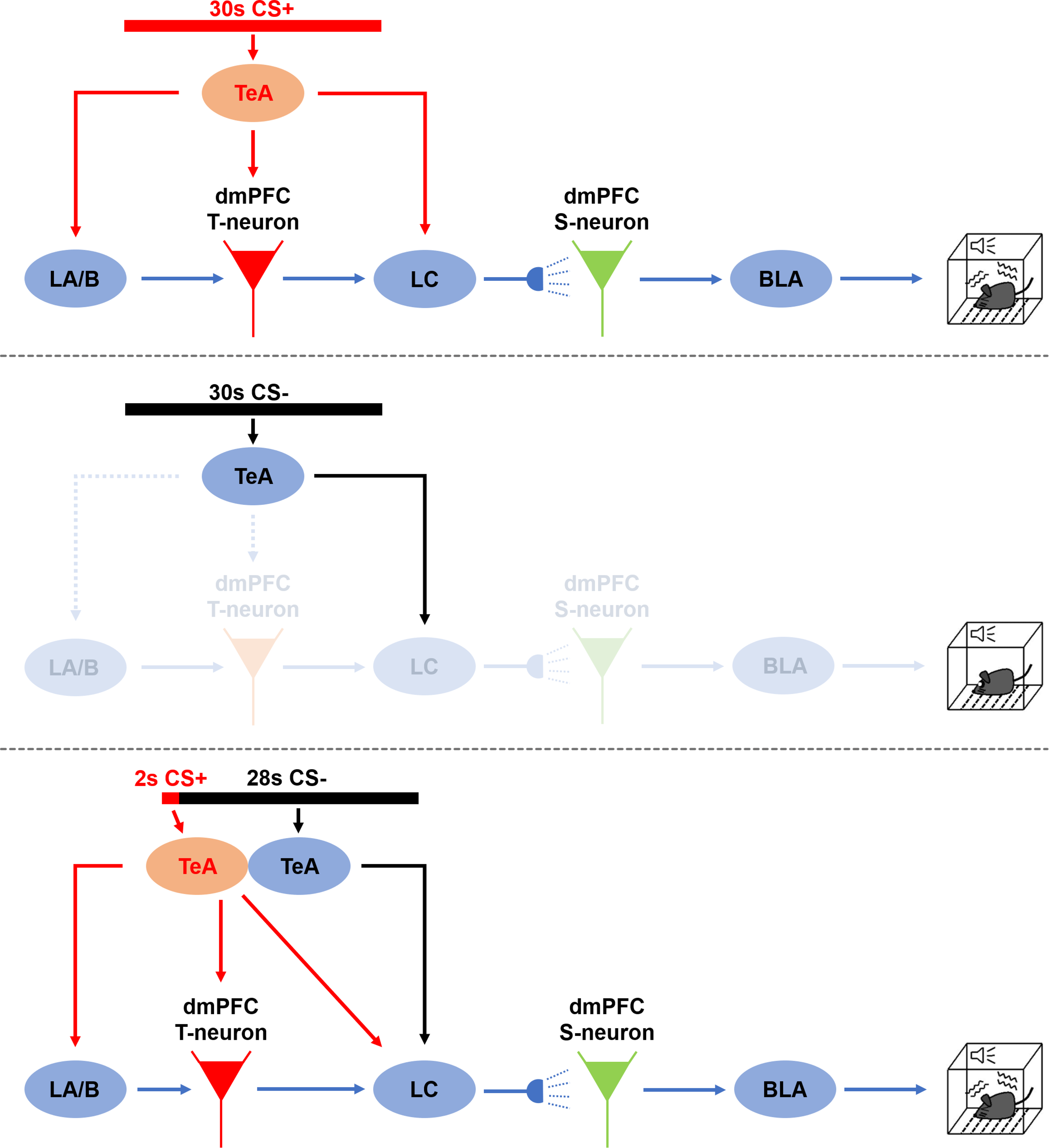
Model of memory to behavior in 3 sets of CSs. (Top, 30 s CS+) CS+ activates TeA, which, in turn, activates LA/B, PFC T-neurons, and LC neurons, S-neurons send outputs to BLA to enable freezing behavior. (Middle, 30 s CS-) CS- only activates outputs to LC, but is unable to activate LC neurons and downstream circuit to elicit freezing behavior. (Bottom, 2 s CS+/28 s CS-) 2 s CS+ activates TeA, which, in turn, activates LA/B, PFC T-neuron, and LC neurons, the ensuing CS- continuously activates LC neurons (similar to previous CS+ that initiates the gating) and downstream circuit to enable freezing behavior.

## Notes

### Competing Interest Statement

The authors have declared no competing interest.

